# Multi-omics subtyping of hepatocellular carcinoma patients using a Bayesian network mixture model

**DOI:** 10.1101/2021.12.16.473083

**Authors:** Polina Suter, Eva Dazert, Jack Kuipers, Charlotte K.Y. Ng, Tuyana Boldanova, Michael N. Hall, Markus H. Heim, Niko Beerenwinkel

## Abstract

Comprehensive molecular characterization of cancer subtypes is essential for predicting clinical outcomes and searching for personalized treatments. We present bnClustOmics, a statistical model and computational tool for multi-omics unsupervised clustering, which serves a dual purpose: Clustering patient samples based on a Bayesian network mixture model and learning the networks of omics variables representing these clusters. The discovered networks encode interactions among all omics variables and provide a molecular characterization of each patient subgroup. We conducted simulation studies that demonstrated the advantages of our approach compared to other clustering methods in the case where the generative model is a mixture of Bayesian networks. We applied bnClustOmics to a hepatocellular carcinoma (HCC) dataset comprising genome (mutation and copy number), transcriptome, proteome, and phosphoproteome data. We identified three main HCC subtypes together with molecular characteristics, some of which are associated with survival even when adjusting for the clinical stage. Cluster-specific networks shed light on the links between genotypes and molecular phenotypes of samples within their respective clusters and suggest targets for personalized treatments.

**Author summary:** Multi-omics approaches to cancer subtyping can provide more insights into molecular changes in tumors compared to single-omics approaches. However, most multi-omics clustering methods do not take into account that gene products interact, for example, as parts of protein complexes or signaling networks. Here we present bnClustOmics, a Bayesian network mixture model for unsupervised clustering of multi-omics data, which can represent dependencies among molecular changes of various omics types explicitly. Unlike other approaches that use data from public interaction databases as ground truth, bnClustOmics learns the dependencies between genes from the analyzed multi-omics dataset. At the same time, our approach can also account for prior knowledge from public interaction databases and use it to guide network learning without losing the ability to learn new dependencies. We applied bnClustOmics to a multi-omics HCC dataset and identified three subtypes similar to those identified in other HCC studies. The cluster-specific networks learned by bnClustOmics revealed additional insights into the molecular characterization of the discovered subgroups and highlighted the changes in signaling networks leading to distinct HCC phenotypes.

## Introduction

Cancer is a complex disease and one of the leading causes of death worldwide. Over the last decades, much research was devoted to discovering cancer subtypes based on genomic and transcriptomic data [1–3]. Molecular subtyping approaches based on gene expression have been helpful for the identification of markers associated with clinical outcomes and facilitated the search for targeted therapies [4,5]. More recently, cancer subtyping has been based on integrating multiple different omics types [6–9]. Multiple tools have been developed to integrate multi-omics data and learn interaction networks to understand what drives oncogenesis [10,11]. However, our understanding of how heterogeneous genetic alterations in cancer cells affect signaling pathways and lead to a few disease phenotypes is still far from complete [12,13]. One major obstacle is the missing connection between methods for network discovery and approaches to molecular subtyping. Almost all existing methods focus on only one of these two tasks.

Only a few multi-omics clustering methods include interactions between gene products into the model explicitly. Some of them are designed for single omics types [14,15] or use a supervised approach for clustering [16]. PARADIGM [17] and BiCON [18] unsupervised clustering (of patient samples) while accounting for the fact that genes products can interact with each other and that interactions may differ between patient groups. However, these methods rely entirely on existing protein-protein interaction (PPI) databases and consider them as ground truth. Instead of learning the network from the dataset, they map the omics data onto interactions from existing databases by considering pairwise dependencies between genes. Such approaches are prone to mistakes contained in databases and do not allow the discovery of unknown interactions.

When learning gene regulatory networks, the Bayesian network framework is often used instead of pairwise correlation analysis since it can uncover direct interactions and, in some cases, learn their directions [19,20]. A Bayesian network mixture model was used for clustering of pan-cancer mutation data [14], but never applied to any other omics types or integrated multi-omics model for unsupervised clustering.

Here, we extend the model of Kuipers et al. [14] to multi-omics data comprising discrete and continuous data types. We present bnClustOmics, an unsupervised clustering method based on the assumption that the cancer subtype can be represented as a Bayesian network consisting of omics variables of various types. Our model reflects the consensus view of cancer mechanisms, in which genetic alterations disrupt normal cell signaling and activate oncogenic pathways. Biological experiments have shown that mutations in cancer cells result in altered interactions between proteins, including phosphoproteins [21]. Thus, modeling the subtype-specific changes in the interactome may improve the clustering model. With cancer subtypes being modeled as Bayesian networks, bnClustOmics can detect the signal from interactions that differ in networks representing different subtypes. A major advantage of bnClustOmics compared to other methods for multi-omics clustering is that the output includes networks (learned *de novo*) representing discovered clusters which can be considered further in downstream analyses and shed light on subtype-specific cancer mechanisms.

We demonstrated in simulation studies that many commonly used clustering methods, including those specifically designed for multi-omics data, have a limited ability to detect a signal from changed interactions, whereas the ability of bnClustOmics to do so improves its clustering accuracy. In particular, we compared traditional clustering approaches with three different approaches designed for multi-omics data. Among multi-omics approaches, we selected methods that demonstrated good performance in previous benchmarking studies [6,22]. iClusterPlus [23] assumes that only a limited number of features are relevant and uses regularization to select features, while CIMLR [24] can incorporate the complete genome without enforcing sparsity. CIMLR was expected to perform better in a broad range of settings due to its claimed ability to learn the importance of different omics types from the analyzed dataset [24], however Duan et al. reported controversial results in this regard [22]. We also added MOFA [25] to benchmarking since it demonstrated good results with regard to feature selection in our simulations. bnClustOmics is only feasible for a limited number of omics features, hence the importance of each omics type is implicitly affected by the feature selection method. We tried several approaches to select relevant features and compared the performance of bnClustOmics using a selected subset to all other clustering methods applied to a non-reduced set.

We applied bnClustOmics to a multi-omics dataset from hepatocellular carcinoma (HCC) patient biopsies [26]. HCC is the most common type of primary liver cancer, which is the fourth most common cause of cancer-related mortality worldwide [27]. We discovered three clusters of HCC patients based on five omics types: mutations and copy number changes (both genome), transcriptome, proteome, and phosphoproteome. The number and molecular characteristics of the three discovered groups confirm many findings from previous HCC studies, including an analysis of the same HCC dataset [26]. In addition to cluster assignments, we analyzed the cluster-specific networks learned by bnClustOmics and scrutinized specific edges which connect changes in the genome to abnormal expression of transcripts, proteins, and phosphorylation sites. Furthermore, we identified hub nodes, i.e. genes with the most stable and most varying neighbors across cluster-specific networks based on the posterior probabilities of the edges. Cluster-specific connections between omics variables provide insights into the molecular characteristics underlying HCC subtypes and suggest targets for personalized therapies.

## 1 Results and Discussion

### 1.1 Model and workflow

We model a cancer subtype as a Bayesian network, whose nodes represent different omics measurements of the same set of genes. The HCC dataset [26] includes five omics types, namely mutations and copy number changes (both genome, denoted *M* and *CN*), transcriptome (*T*), proteome (*P*), and phosphoproteome (*PP*). The edges in the network represent statistical dependencies among all observations across all omics types. Such dependencies are not limited to a single biological interpretation. For example, an edge in the network might represent a physical interaction between proteins, a regulatory relationship between a transcription factor and its target, a functional interaction or a co-expression pattern. A functional interaction denotes an indirect association where two gene products do not physically interact but are jointly involved in the same cellular process [28].

By design, our integrative model prohibits any edges from nodes of continuous data types (*T*, *P*, *PP*) to nodes of binary or ordinal data types (*M*, *CN*). Prohibiting interactions between certain omics types avoids overfitting and results in more interpretable networks. We only allow edges aligned with the information flow of the central dogma of molecular biology [29].

At the first step of the analysis, we perform feature selection from all features of all available omics types (Fig 1). To analyze the HCC dataset, we selected features based on multi-omics factor analysis (MOFA, [25]) latent factor analysis, differential gene expression (DGE) analysis, and prior knowledge about signaling networks (Section 2.13).

**Fig 1.**
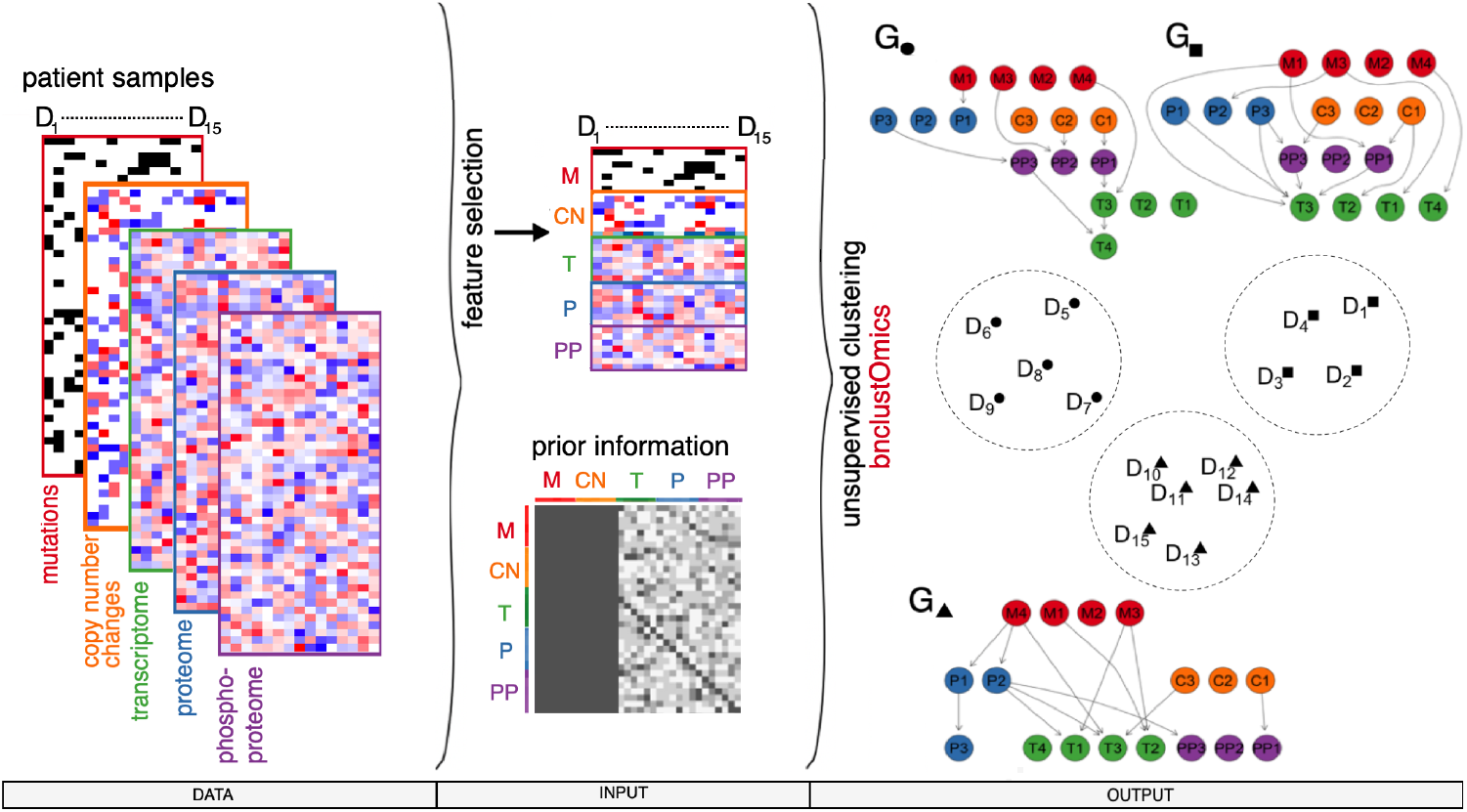
Bayesian network-based clustering workflow. Multiple omics types, both binary and continuous, are allowed as input data types (left). After feature selection is performed, prior knowledge about interactions between nodes can be included via blacklisting and penalization matrices (middle). bnClustOmics performs unsupervised clustering based on the selected features, blacklist, and penalization matrices. The output (right) includes cluster assignments (encircled patient sample), cluster-specific networks, and posterior probabilities of all individual edges in these graphs. Here, three patient clusters are depicted and labeled ●, ▲, and ■.

bnClustOmics uses a Bayesian network mixture model and employs the EM algorithm [14] to cluster patient samples and learn the networks representing those clusters. Unlike other multi-omics clustering methods, bnClustOmics does not rely on interactions from databases, but learns Bayesian networks from data *de novo* using a Bayesian approach [14,30,31]. However, it is possible to construct blacklist and penalization matrices that incorporate prior information about interactions between selected features and guide network learning in subsequent steps. In the extreme case, we could blacklist all edges which are not found in a specific database. However, blacklisting prevents the discovery of new interactions. Instead, we can use edge-specific penalization factors to modify the prior probability distribution of the graph structure and lower the probability of such edges appearing in the resulting graphs. The penalization matrix also provides an easy way to incorporate a confidence score which is often assigned to interactions in the PPI databases.

bnClustOmics takes as input the observed values of the selected omics features for all patients, the number of clusters, and optional blacklist and penalization matrices. As output, we obtain cluster assignments for all patient samples, cluster-specific networks consisting of omics variables, and the log-likelihood, AIC and BIC scores of the estimated model. The AIC and BIC can be used to determine the optimal number of clusters. In addition, the Bayesian method used for structure learning provides estimates of posterior probabilities of all edges in the discovered networks. The statistical model presented in this work is implemented in the R package bnClustOmics and available at the GitHub repository https://github.com/cbg-ethz/bnClustOmics.

### 1.2 Benchmarking

We compared the performance of several clustering algorithms to bnClustOmics when the data generating model is a mixture of Bayesian networks. For comparison, we selected several general clustering methods as well as methods specifically designed for integration and clustering of multi-omics data, including kmeans [32], hclust [32], mclust [33], iClusterPlus [23], CIMLR [24], and MOFA [25].

For each set of simulation settings, we generated 30 – 50 Bayesian network mixtures (Section 2.2), where each directed acyclic graph (DAG) *G_k_* consists of *n_c_* Gaussian and *n_b_* Bernoulli random variables (S1 Appendix). We used the adjusted Rand index (ARI, [34]) between estimated cluster assignments and the ground truth membership as a measure of accuracy.

For large sample sizes, bnClustOmics reaches a high accuracy even in a setting where the difference between cluster centers is small (Fig 2A), while the other algorithms fail to discover cluster assignments when cluster centers are very close to each other. Accuracy improves when the distances between centers of mixture components become larger for all algorithms except CIMLR. In our simulation settings, CIMLR failed to detect the signal from the continuous nodes in the presence of binary nodes. When we removed binary nodes from the simulated datasets and applied CIMLR to the continuous part only, its accuracy improved considerably.

**Fig 2.**
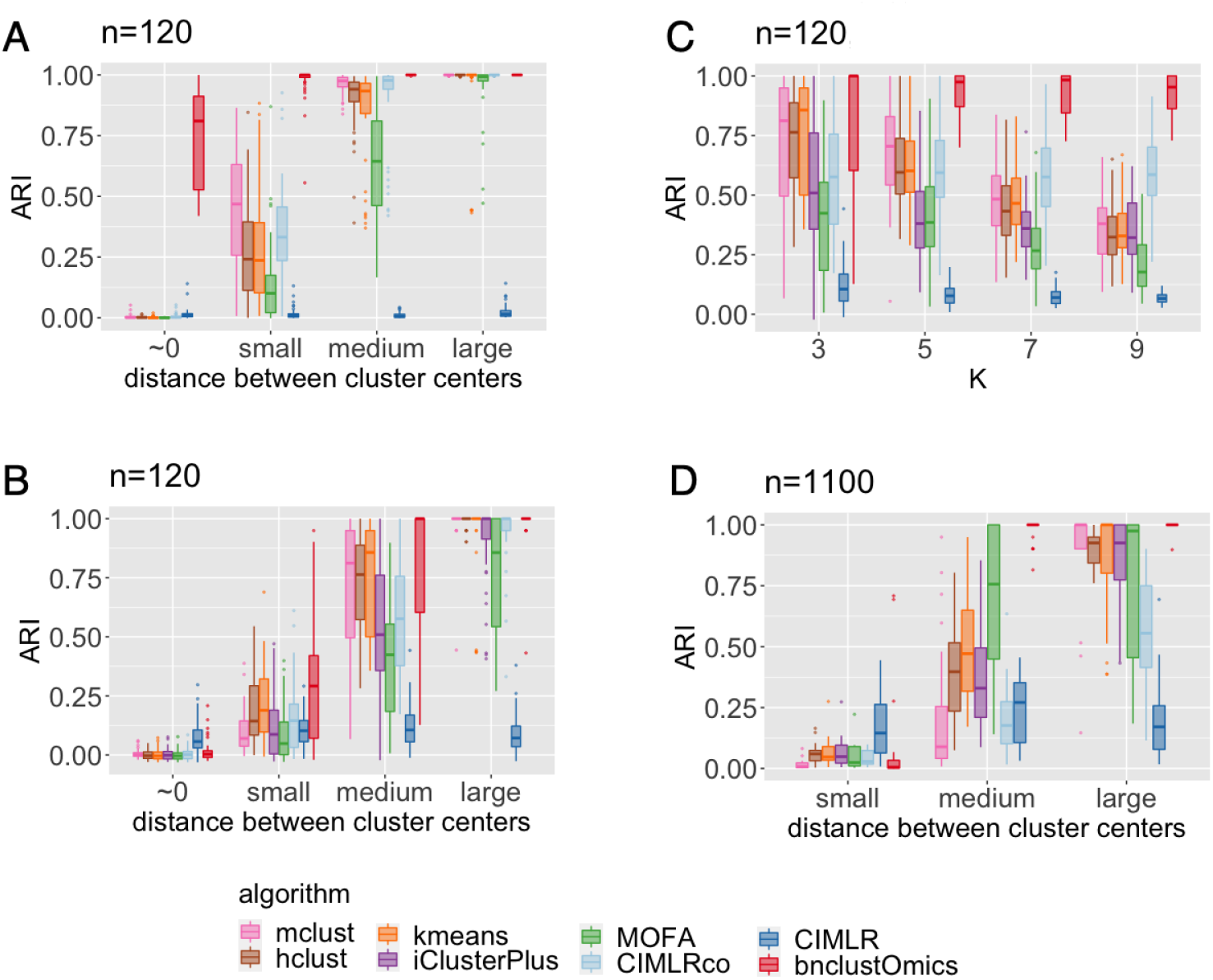
Benchmarking of algorithms for unsupervised clustering of multi-omics data. 50 Bayesian network mixtures were generated for each simulation setting. For general clustering approaches, the dimension was reduced by applying PCA and running clustering on the first 5 principal components. All integrative multi-omics approaches were applied to the original data unless specified otherwise. CIMLRco denotes clustering results of the application of CIMLR to a subset of data consisting of observations of only continuous variables. *N_Z_k__* denotes the number of observations in one cluster, *K* the number of clusters, *n_c_* number of continuous nodes, *n_b_* number of binary nodes in networks. (A) *K* = 3, *n_c_* = 100, *n_b_* = 20, *N_Z_k__* = 200 (B) *K* = 3, *n_c_* = 100, *n_b_* = 20, *N_Z_k__* = 20 (C) *n_c_* = 100, *n_b_* = 20, *N_Z_k__* =20, *K* ∈ {3, 5, 7, 9}, distance between centers set to medium (D) *K* = 3, *n_c_* = 1000, *n_b_* = 100, *N_Z_k__* = 20 bnClustOmics was applied to a subset of data consisting of all binary nodes with non-zero standard deviation and 150 selected continuous nodes.

For small sample sizes, all methods demonstrate lower clustering accuracy (Fig 2B), and bnClustOmics outperforms the other approaches in the majority of cases. We attribute this outperformance to the ability of bnClustOmics to detect the signal not only from differences between cluster centers but also structural differences between graphs representing clusters.

Next, we fix the distance between the centers of the distributions at a medium value and analyze the performance of different algorithms with four different values *K* of the number of clusters. The clustering accuracy of bnClustOmics does not become worse with increasing number of clusters *K*, while for the other algorithms, the accuracy decreases (Fig 2C). Among the other algorithms, CIMLR applied to only a continuous part of the data performs the best. However, its accuracy is again substantially worse when the binary data is included.

Since bnClustOmics is only computationally feasible for networks with a limited number of nodes, its performance may strongly depend on the selection of the relevant features. To assess whether reducing the number of features affects the clustering accuracy of bnClustOmics, we generated Bayesian network mixtures with *n_c_* = 1000 Gaussian nodes and *n_b_* = 100 binary nodes from *K* = 3 mixture components and applied all clustering approaches to this dataset. All algorithms were applied to the complete dataset, except bnClustOmics which was applied to only a subset of the data consisting of 150 continuous features (S2 Appendix) and all binary features with at least one non-zero observation. We found that despite using significantly fewer variables, bnClustOmics outperforms the other methods (Fig 2D) when the distances between cluster centers are medium to large. For small distances, all methods perform poorly.

So far, we assumed that the number of clusters is known; however, in a fully unsupervised setting, this is not the case. bnClustOmics allows estimating the number of clusters *K* using either the AIC or BIC score. Our simulations indicate that for small sample size, AIC works better (Fig 3A), while for large sample size, BIC shows better results (Fig 3B).

**Fig 3.**
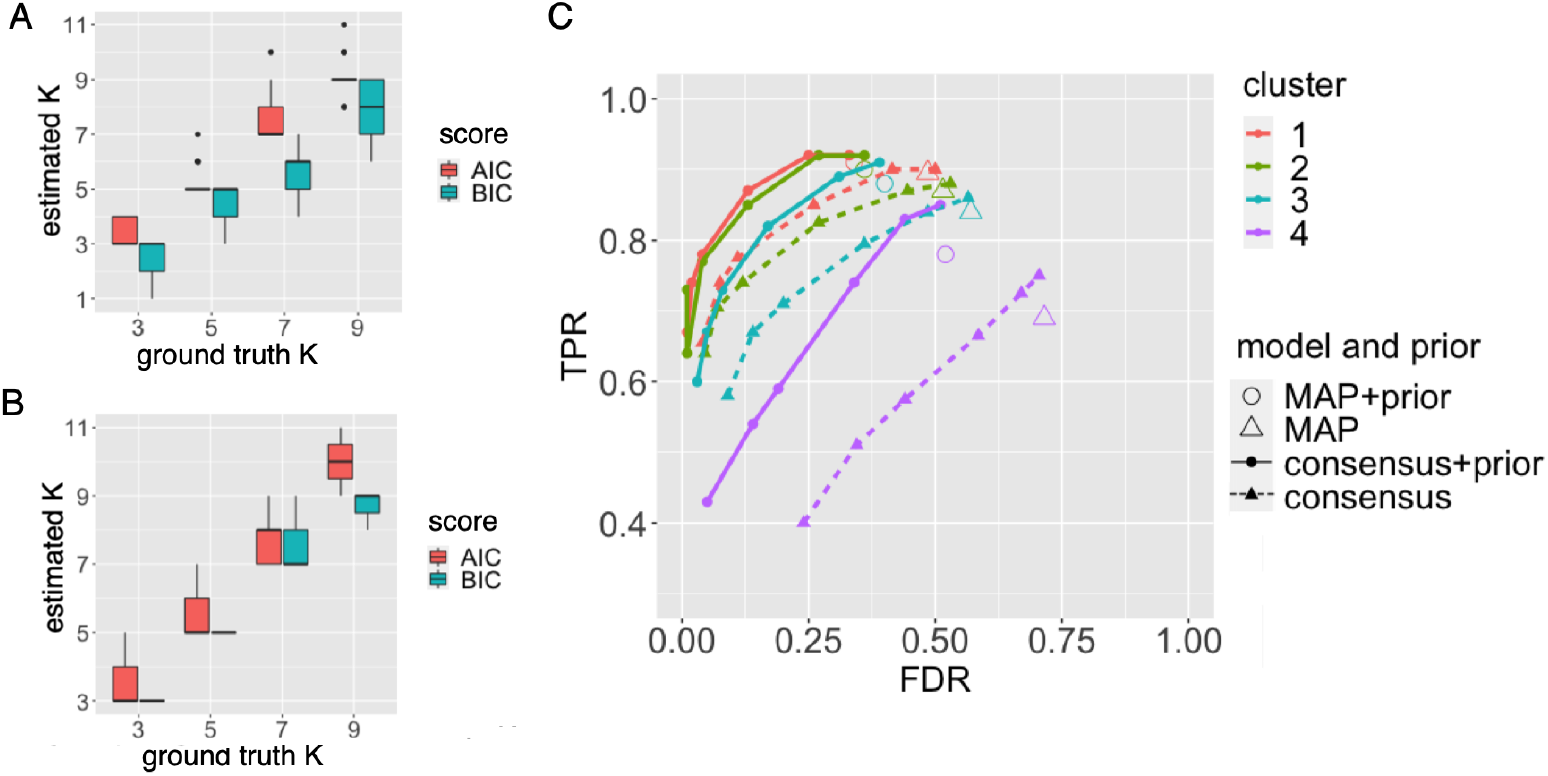
Defining the optimal number of clusters and structure fit. 30 Bayesian network mixtures were generated for each number of clusters *K* ∈ {3, 5, 7, 9} (ground truth). bnClustOmics was applied for each estimated *K* ∈ {1, …, 11} to each generated dataset and 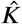 was determined by minimizing the AIC or BIC score. (A) *N_Z_k__* =20 (B) *N_Z_k__* = 200 (C) 50 datasets were generated from Bayesian network mixtures consisting of *K* = 4 components with number of observations *N_Z_k__* ∈ {150, 100, 50, 20} corresponding to cluster 1 (red), cluster 2 (green), cluster 3 (turquoise) and cluster 4 (violet). To construct the penalization matrix (prior), we first defined the edges representing interactions from databases by taking the union of all edges in the ground truth structures. Afterward, we removed 5% of these edges, modeling false-negative interactions in databases, and added 5% of false positives. The entries of the penalization matrix corresponding to the defined set were not penalized; all other edges were penalized by a factor of two. The simulated datasets were clustered using bnClustOmics with and without the penalization matrix. Resulting MAP and consensus models corresponding to posterior thresholds of *p* ∈ {0.3, 0.5, 0.7, 0.9, 0.95, 0.99} were assessed using TPR and FDR.

Next, we tested the ability of bnClustOmics to reconstruct Bayesian networks representing discovered clusters. We generated 50 Bayesian network mixtures with *K* = 4 components and unequal weights, such that the four clusters contain 150, 100, 50, and 20 observations, respectively. The Bayesian approach yields estimated *maximum a posteriori* (MAP) structures, i.e. graphs which have the highest scores of all considered structures and represent the best fit to the data. In addition to MAP graphs, we also estimated the consensus structures (Section 2.5), which consist of edges whose posterior probabilities are higher than a certain threshold.

The number of observations per cluster correlates positively with the accuracy of the learned MAP structures, as progressively higher TPR and lower FDR levels were reported for MAP structures corresponding to a higher number of observations (Fig 3C). However, the FDR of MAP structures is rather high, especially for the cluster 4 with the smallest number of observations. We observe that consensus graphs help reduce FDR compared to the MAP estimates, although at the cost of reducing the true positive rate (TPR). The structural Hamming distance (SHD) is smaller for consensus structures than for MAP structures (S1 Fig). In our simulation, a posterior threshold of 0.7 minimizes the SHD for *N_Z_k__* ∈ {150, 100, 50} and 0.95 for *N_Z_k__* = 20.

The Bayesian approach allows us to include prior knowledge about known interactions and guide *de novo* network learning. In the analysis of mutation data, an edge penalization matrix was used by Kuipers et al. [14] to include prior information from the database STRING [35]. The edge penalization matrix is used to modify the prior over structures, such that penalized edges have a lower chance to appear in the discovered graphs (Section 2.9). PPI databases contain known interactions between genes but most often do not describe the context in which a particular interaction occurs. Hence, if interactions differ between unknown cancer subtypes, we cannot learn them using a database alone. To assess to which extent the penalization matrix can improve network discovery, we constructed a simulated database of interactions by taking the union of all edges in the ground truth structures and introducing 10% of mistakes which model false-positive (5%) and false-negative (5%) interactions in databases. The entries of the penalization matrix corresponding to interactions from the simulated database were not penalized; all other edges were penalized by a factor of two.

The usage of an edge penalization matrix resulted in MAP and consensus structures containing fewer false-positive edges and more true positives than corresponding structures obtained without using a penalization matrix (Fig 3C). Limited sample size is a common problem of biological data, and proteome and phosphoproteome data are generally scarce. At the same time, extensive databases exist which include known protein-protein interactions and regulatory relationships identified in biological or computational studies. Hence, including information from such databases can be helpful for network reconstruction.

### 1.3 HCC patient subtyping

We analyzed the HCC multi-omics dataset [26] comprising 50 biopsies from 48 patients and including five omics types, namely mutations and CNAs (both genome), transcriptome, proteome, and phosphoproteome. In order to apply bnClustOmics, we first performed feature selection as follows. To select *M* features, we used a list of significantly mutated genes in the analyzed cohort identified by Ng et al. [26]. In addition, we included possible drivers of HCC identified in other studies [36–38]. To select continuous features, we applied MOFA and performed latent factor analysis. In addition, we included the *P* and *PP* features, which are differentially expressed/phosphorylated in tumor samples and either are present in the kinase-substrate database, or are known transcription factors according to the Omnipath database [39] (Section 2.10). We proceeded with the construction of the blacklist and penalization matrices as described in Section 2.8 and Section 2.9 and included prior information about interactions between selected features from the STRING and Omnipath databases [35,39].

We ran the algorithm for *K* =1, 2, 3, 4, and 5 clusters. The BIC and AIC scores indicated *K* = 3 as the optimal number of clusters (Fig 4A). *K* = 3 clusters were also found as optimal for the same data in [26] and in another HCC study applying a network-based method to the TCGA HCC dataset [40]. Similar to the clusters discovered in [26], the clusters discovered by bnClustOmics (Fig 4D) are associated with mutations in the genes *TP53* and *CTNNB1*, Edmondson grade, and BCLC stage (*p*-values using Fisher’s exact test are 0.012, 0.001, 0.007, and 0.019, respectively). Cluster 1 is dominated by samples with mutations in *CTNNB1*, and cluster 2 is dominated by samples with mutations in *TP53* (Fig 4C). Cluster 3 is the most heterogeneous in terms of mutations. However, all 4 samples with mutations in *ALB* are in cluster 3.

**Fig 4.**
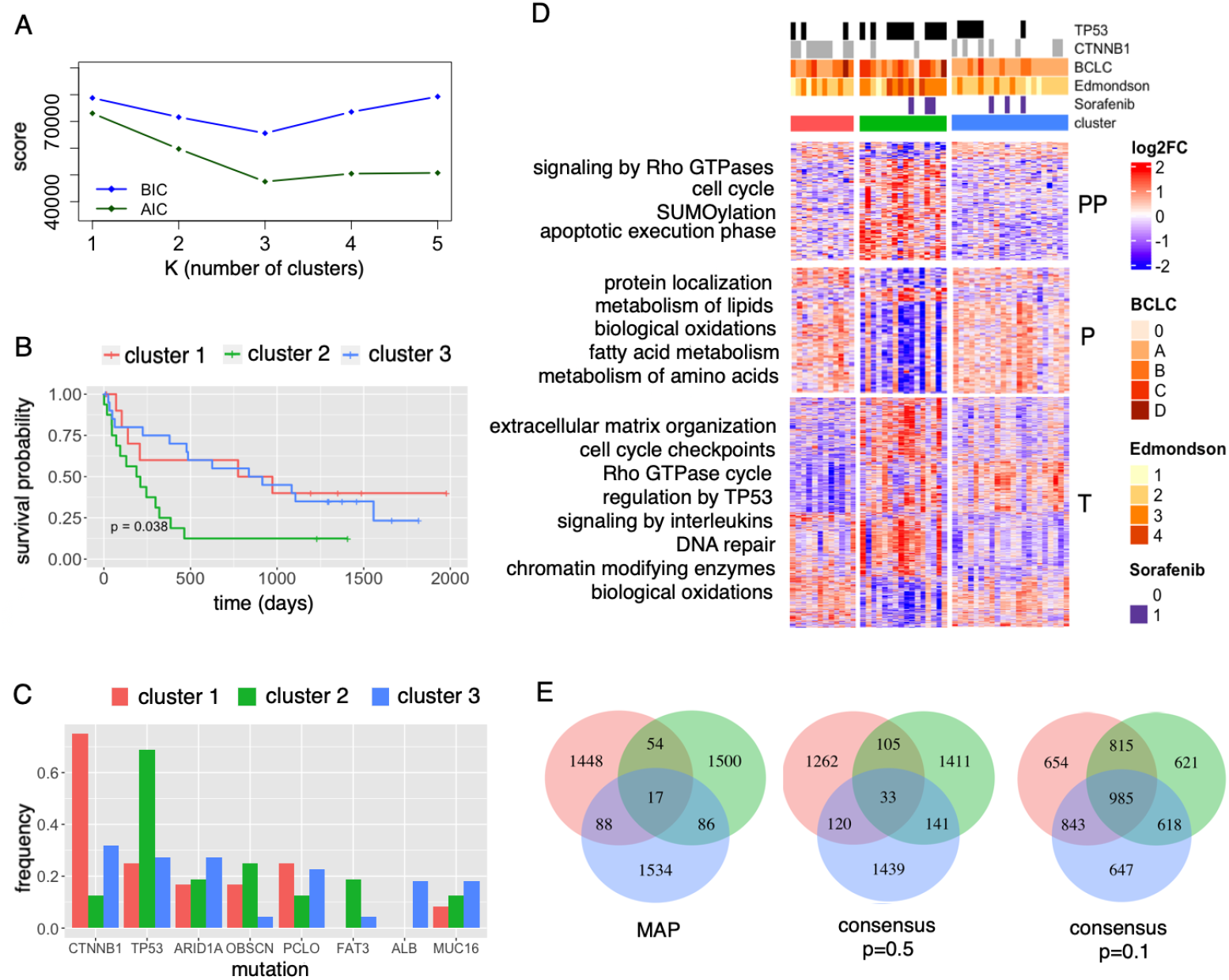
Multi-omics clustering of the HCC dataset with bnClustOmics. (A) BIC and AIC scores of models with different numbers of clusters. (B) Kaplan-Meier survival curves for patients in discovered clusters. (C) Mutational frequencies in discovered clusters. Only mutations with frequency ≥ 15% in at least one of the clusters are shown. (D) Pathway enrichment differences between clusters. (E) Venn diagrams showing the number of common and cluster-specific edges in the discovered MAP and consensus networks learned for cluster 1 (red), cluster 2 (green), cluster 3 (blue); edge directions were disregarded.

The three discovered subgroups are associated with patient survival with and without adjustment for BCLC stage (S2 Fig, Section 2.11). In particular, the Cox proportional hazards model revealed that cluster 2 is associated with a poor prognosis (*p* = 0.039 for the non-adjusted model and *p* = 0.024 for the adjusted model), while survival prognoses for cluster 1 and cluster 3 are better and similar. We tested several other approaches for multi-omics clustering, including MOFA, which we used for feature selection. None of the models produced patient subgroups significantly associated with survival when adjusting for BCLC stage (S1 Table, Survival analysis).

To identify processes whose regulation is different between the three patient clusters, we performed DGE and pathway enrichment analysis (Section 2.12). The differences in enriched pathways at all omics levels are more pronounced between cluster 2 and the other clusters (Fig 4D). Significant differences in enriched pathways between cluster 1 and cluster 3 were identified only at the transcriptome level, but not the proteome or phosphoproteome level. However, this situation can result from a combination of noisy data and limitations of pathway enrichment analysis [41].

In order to extend the molecular characterization of the discovered clusters beyond expression levels and mutational frequencies, we analyzed the multi-omics networks that define the clusters. The three MAP networks are very different from each other (Fig 4E). At the same time, the similarities between consensus networks constructed at the edge-wise posterior level of 0.1 are substantially larger (Section 2.5). While 0.1 is a low confidence threshold, the proportion of edges that pass this threshold is around 2% of all non-blacklisted edges for each network. Therefore, the high degree of similarity at the 0.1 level suggests that the posterior landscapes are not as different as the MAP structures. This reflects a high level of modeling uncertainty due to the small effective sample sizes from which the networks were learned. The downside of MAP structures is the inability to account for this uncertainty which can lead to overfitting, as we have seen in simulation studies (Fig 3C).

To address this limitation, we took advantage of the Bayesian approach that we used for structure learning and using several posterior thresholds constructed consensus networks for downstream analysis (Section 2.5).

### 1.4 Downstream effects of mutated genes

bnClustOmics allows for identifying links between genotypes and molecular characteristics of individual clusters. We analyzed all children of *M* (mutation) nodes in the cluster-specific networks. At the first step, we performed pathway enrichment analysis and identified KEGG pathways that are enriched with cluster-specific children of mutation nodes (S2 Table). Signaling pathways associated with HCC, including PIK3-Akt, p53 and cell cycle, were enriched in all clusters. The differences in enriched pathways between the clusters can be connected to their genotypes. For example, the Wnt signaling pathway, whose activation is usually associated with mutations in *CTNNB1* is enriched in cluster 1 and cluster 3 but not in cluster 2. Network *G*_3_ is characterized by more connections than other networks due to a higher level of heterogeneity in cluster 3. As a result, more pathways were found to be enriched with direct neighbors of *M* nodes.

We further scrutinized the individual edges connecting mutations to the nodes representing genes products of other omics types. To find the connections which can explain the abnormal expression of *T*, *P* and *PP* nodes, we have selected children of all mutation nodes in all networks which are differentially expressed in at least one cluster or in the whole dataset (Fig 5). The frequently mutated genes *TP53, CTNNB1*, and *ARID1A* have the most children across networks. However, *CTNNB1* has the largest proportion of children that are the same across the clusters, whereas the children of *ARID1A* are rather different across the clusters. This suggests that the effects of mutations in *CTNNB1* are more homogeneous, while effects of *ARID1A* are more heterogeneous across clusters. *ARID1A* is a sub-unit of chromatin remodeling complex SWI/SNF and may have broad effects on gene expression levels. Heterogeneous roles of *ARID1A* in HCC were already pointed out in previous studies [42]. In particular, *ARID1A* was found to act both as a tumor suppressor and oncogene depending on the context.

**Fig 5.**
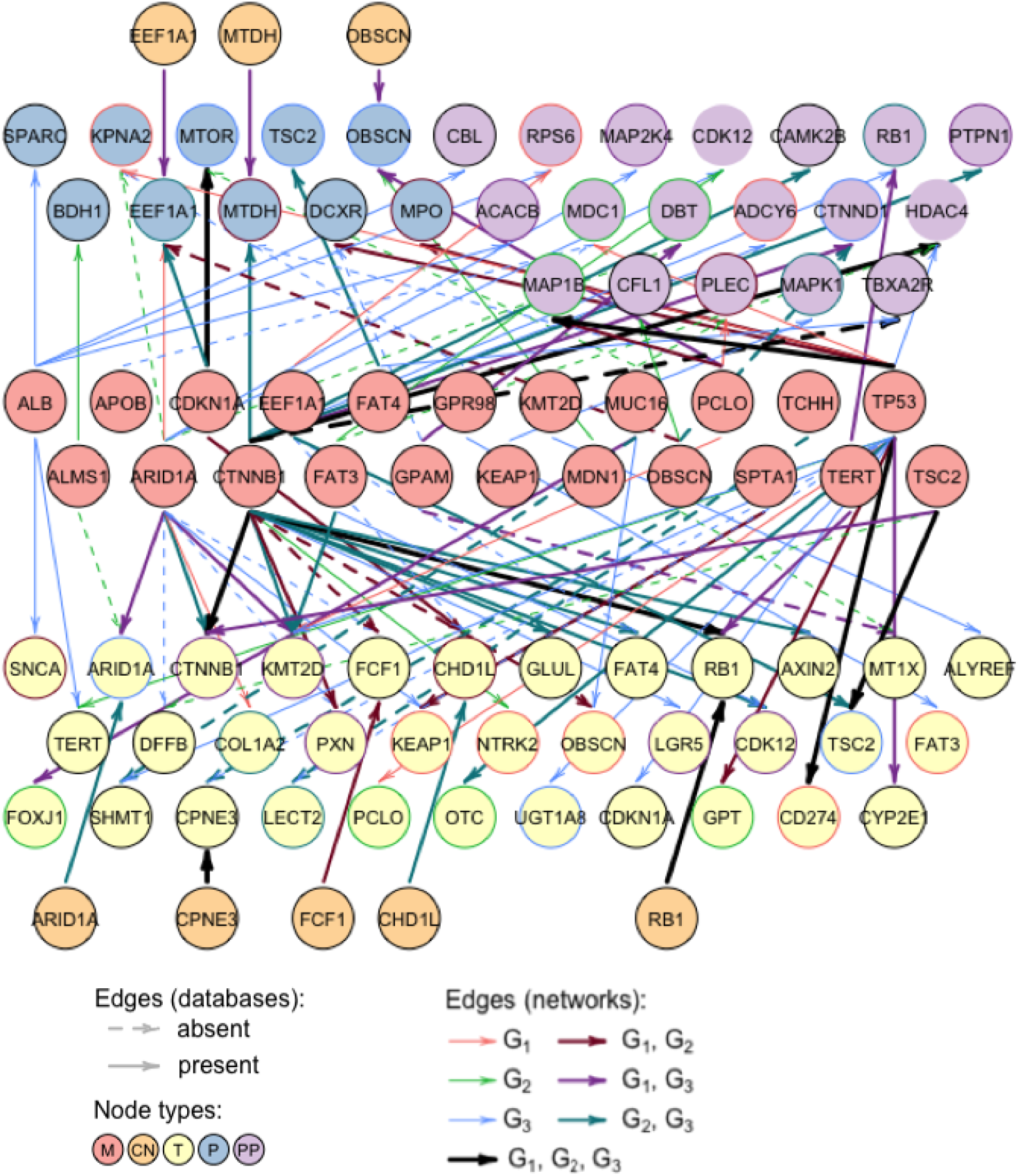
Mutated genes and their most common interaction partners in HCC networks learned by bnClustOmics. Only those *T*, *P*, and *PP* nodes are shown that are differentially expressed/phosphorylated in at least one cluster or the whole dataset. Edges are shown based on their posterior probability: either if they have a high total posterior probability (sum across clusters is at least 1.2), or if they have a high posterior probability in at least one of the clusters (*p* > 0.9). Edge colors indicate in which cluster-specific networks the edges are present with a posterior probability *p* > 0.4: red(*G*_1_), green(*G*_2_), blue (*G*_3_), brown (*G*_1_ and *G*_2_), violet (*G*_1_ and *G*_3_), turquoise (*G*_2_ and *G*_3_), black (*G*_1_ and *G*_2_ and *G*_3_). Border colors of *T*, *P*, and *PP* nodes represent the differential expression status (color scheme is the same as edge colors). Solid edges denote either connections between two omics types of the same gene or interactions found in the STRING and Omnipath databases.

We noted that bnClustOmics was able to capture some of the well-known HCC-specific interactions while performing *de novo* clustering. One example of homogeneous connections is the edge from mutation in *CTNNB1* (denoted *CTNNB1-M*) to the *CTNNB1* transcript abundance (*CTNNB1-T*). In all clusters, the mutation status of *CTNNB1* is positively correlated with the expression of the *CTNNB1* transcript. In cluster 1 and cluster 3, *CTNNB1-T* is overexpressed compared to normal samples. This corresponds to the known effects of *CTNNB1* mutations in HCC [43]. However, in cluster 2, *CTNNB1-T* is not overexpressed, despite the edge between *CTNNB1-M* and *CTNNB1-T*. This situation results from cluster 2 containing only two samples with mutated *CTNNB1* and the fact that mutations in *TP53* are not associated with increased *CTNNB1-T*. This example demonstrates the complementary roles of network analysis with DGE in the downstream analysis.

The edge from *CTNNB1-M* to *GLUL-T* which is present in *G*_2_ and *G*_3_ is another example of a previously known interaction. *GLUL* is known to be upregulated in HCC and is associated with the mutated *CTNNB1*. It is also known that *GLUL* is affected by activation of the Wnt/*β*-catenin pathway at the transcription level, so the incoming edges in the *GLUL-T* node are consistent with previous findings [44]. Interestingly, there is no edge connecting *CTNNB1-M* and *GLUL-T* in *G*_1_. If we examine the interaction partners of *GLUL-T* (Fig 6A), there is an incoming edge that is specific to *G*_1_ coming from the phosphorylation site AXIN2_S70, and AXIN2_S70 has an incoming edge from *CTNNB1* also only in *G*_1_. AXIN2, just like *GLUL*, is a known target of the Wnt/*β*-catenin pathway [45]. The link between proteins GLUL and AXIN2 is also present in the STRING database with an interaction score of 0.42. The phosphorylation site AXIN2_S70 has been mentioned in the study connecting mutations to signaling in breast cancer [46]; however, there have been no previous studies about this phosphorylation site in HCC. Thus, the different path from *CTNNB1-M* to *GLUL* in *G*_1_ compared to *G*_2_ and *G*_3_ may represent differences in signaling leading to the same target. Alternatively, due to a limited number of observations, we may have captured the same process with a different set of edges, so further experiments are needed to clarify this link.

**Fig 6.**
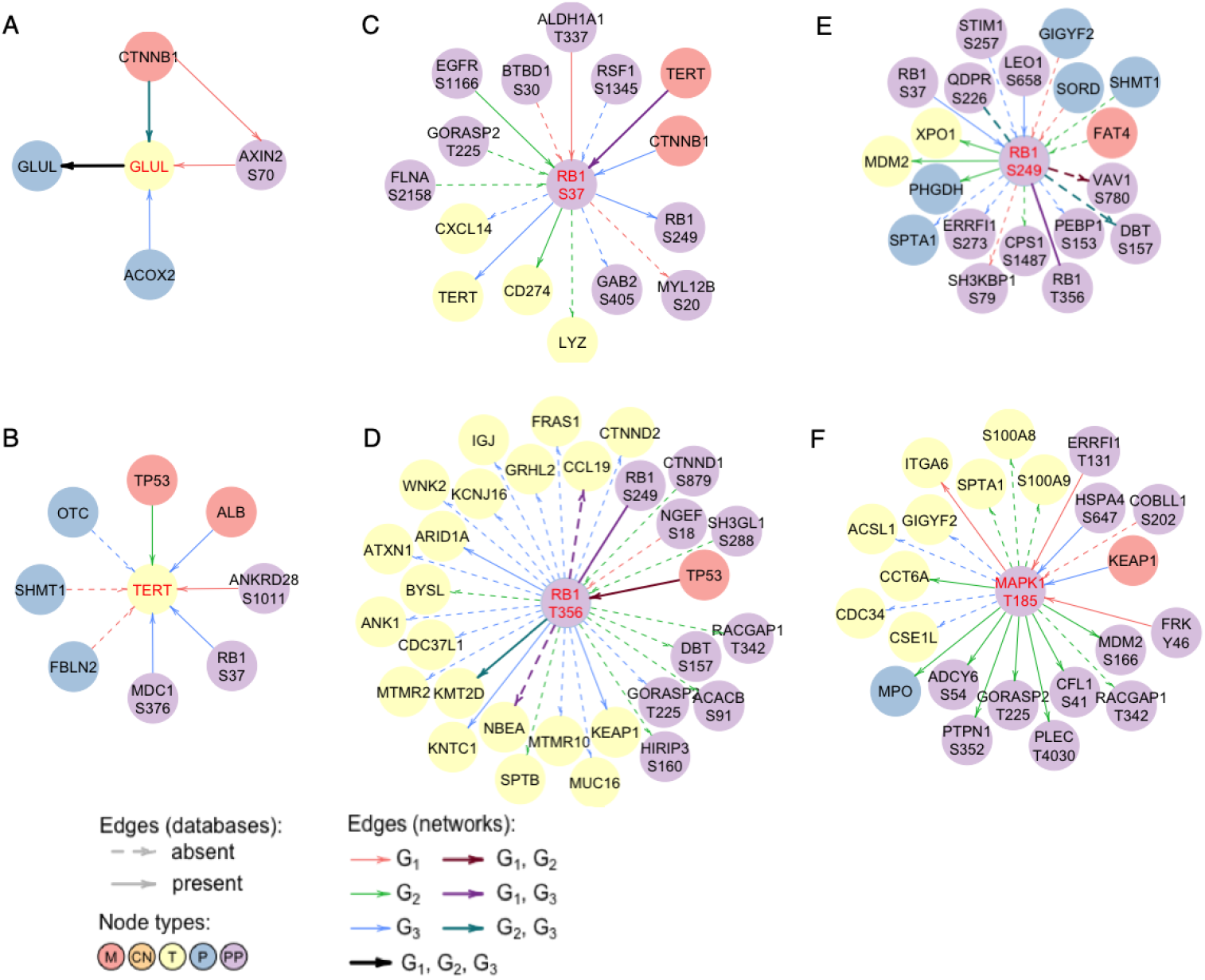
Neighborhoods of individual nodes in the networks learned by bnClustOmics. Direct neighbors of nodes (A) *GLUL-T* (B) *TERT-T* (C) RB1-S37 (D) RB1_T356 (E) RB1-S249 (F) MAPK1_T185 in multi-omics networks discovered by bnClustOmics. Interactions are only shown between the central node and all of its direct neighbors with exception of (A) where we also show the connection between *CTNNB1-M* and AXIN2_S70.

In addition to edges corresponding to known interaction contexts, bnClustOmics discovered edges pointing at new context-specific dependencies. Cluster 2 is characterized by mutations in the *TP53* gene, and we analyzed *TP53-M* connections which might contribute to the phenotype of cluster 2 (S8 Appendix). The transcript node *TERT-T* is differentially expressed in cluster 2 and also has an incoming edge from *TP53-M* in *G*_2_. *TERT-T* expression is known to be upregulated in many cancers including HCC [47] and it is also significantly overexpressed in all clusters in the analyzed cohort. However, the expression level of *TERT-T* is significantly higher in cluster 2 than in cluster 1 and cluster 3 (Figure 9 in S8 Appendix). The high degree of *TERT-T* overexpression is associated with mutations in *TP53* as suggested by *G*_2_. At the same time, the edge from *TP53-M* to *TERT-T* is absent in *G*_1_ and *G*_3_ (Figure 8 in S8 Appendix), suggesting that the effect of mutated *TP53* on *TERT-T* is only present in cluster 2.

**Fig 7.**
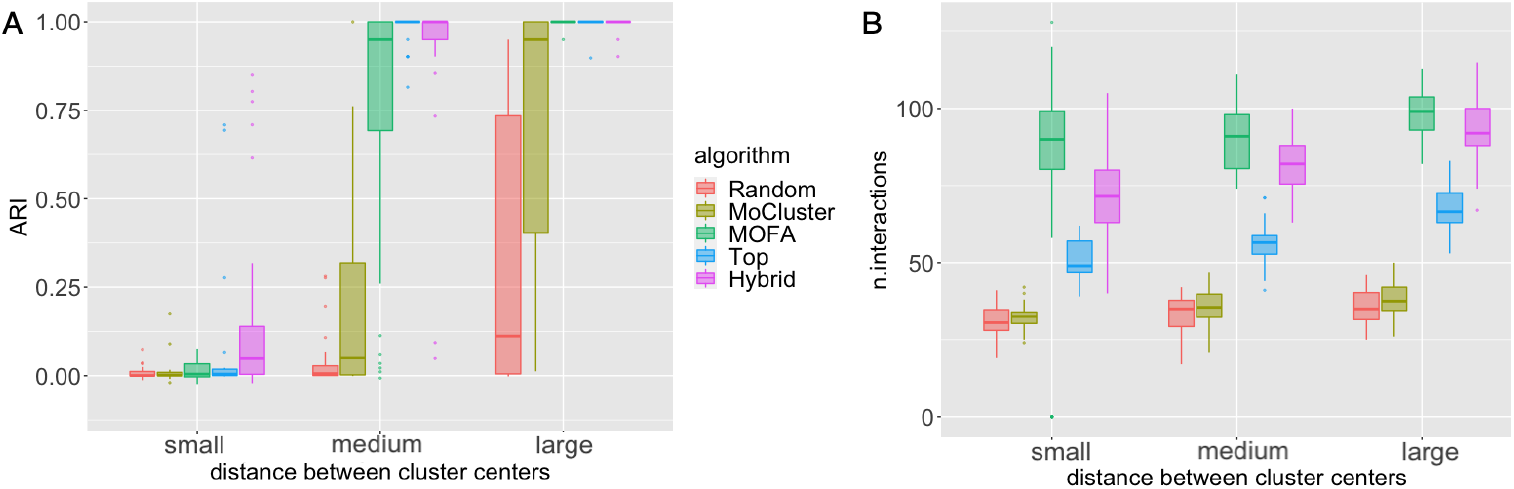
(A) Accuracy of bnClustOmics with different feature selection approaches. (B) The total number of edges from all generated networks which are also present in subnetworks consisting of selected nodes.

**Fig 8.**
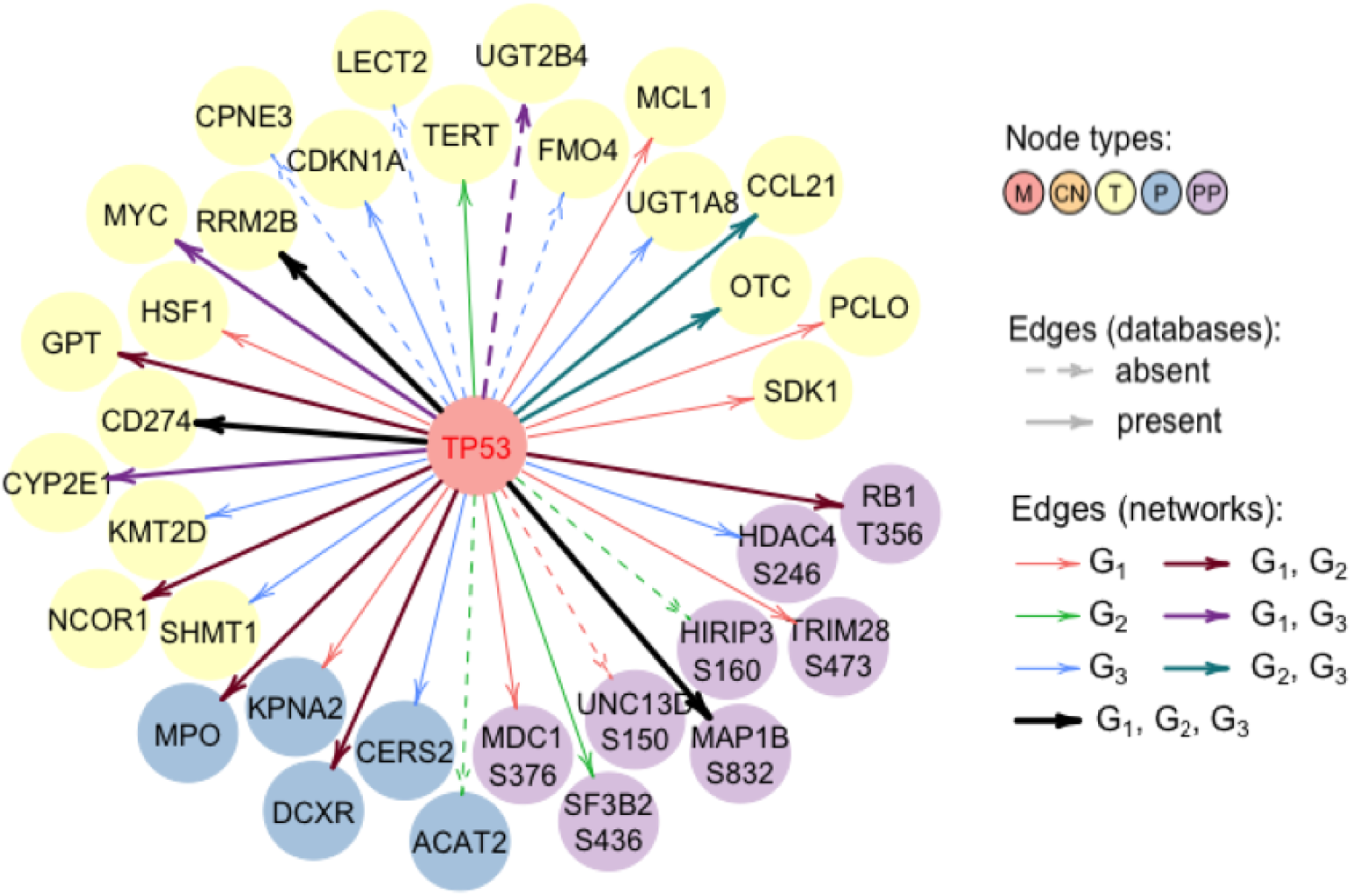
*TP53-M* node and its neighbors in networks representing three clusters identified by bnClustOmics.

**Fig 9.**
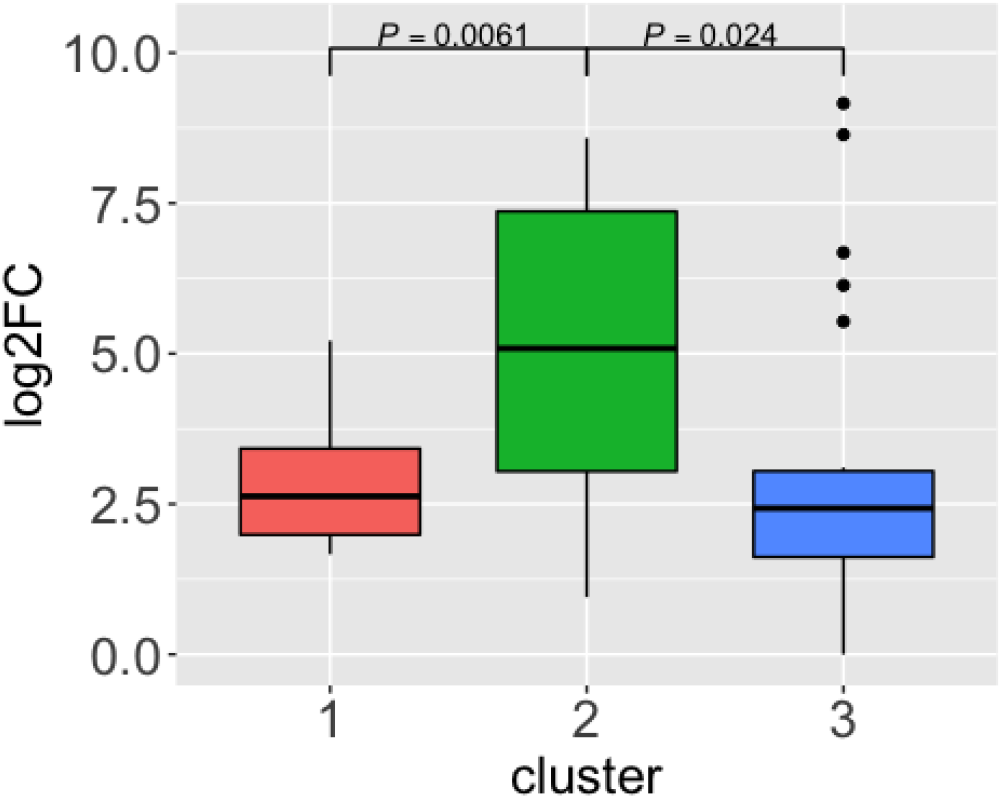
Log2-fold changes between expression of *TERT-T* in three HCC clusters and mean expression of *TERT-T* in 15 healthy livers.

We investigated connections of *TERT-T* in other networks to identify possible sources of its upregulation in remaining clusters (Fig 6B). There is an incoming edge from ALB-*M* in *G*_3_; however, it is negatively correlated with *TERT-T* expression, so mutations in ALB-*M* do not seem to contribute to *TERT-T* overexpression. In addition, there is a *G*_3_-specific incoming edge from the phosphorylation site RB1_S37, which is overexpressed in cluster 2 and cluster 3, but not in cluster 1. Network *G*_3_ suggests that RB1_S37 is associated with overexpression of *TERT-T* and hence might also contribute to carcinogenesis. In *G*_3_, there is an incoming edge to RB1_S37 from *CTNNB1-M* and the corresponding correlation is positive. This suggests that *CTNNB1-M* contributes to RB1_S37 overexpression and via RB1_S37 may affect *TERT-T* as well. However, this dependency is not direct and weaker than the edge from *TP53-M* to *TERT-T* in cluster 2, suggesting that the direction of the effect of mutations in *CTNNB1* and *TP53* on the expression of *TERT* is the same, but the effect size is different. This finding aligns with the associations between mutations in *CTNNB1* and *TP53* and survival. Both mutated genes are drivers of HCC however, *TP53* results in a poorer prognosis than *CTNNB1*.

Other RB1 phosphorylation sites, namely S249 and T356, are highly phosphorylated across all clusters. Moreover, we observe several incoming edges from *M* nodes in all RB1 phosphorylation sites (Fig 6C:E). The mutation statuses of parent nodes of RB1 (*FAT4-M* in cluster 2, *TERT-M* in cluster 3, *TP53-M* in cluster 1) are positively correlated with increased phosphorylation of the respective sites, suggesting that they all may contribute to RB1 hyperphosphorylation. In previous studies, RB1 has been shown to play an important but complex role in cell cycle regulation and apoptosis [48]. It can act both as a tumor suppressor and oncogene depending on its phosphorylation status. All three phosphorylation sites included in our network can be found in the PhosphoSitePlus database [49]. The role of S249 and T356 phosphorylation is well studied and known to affect the cell cycle and apoptosis. The role of S37 phosphorylation is less well known, and there are no studies about its role in HCC. As previously noted, our analysis suggests that phosphorylation of this site may also play a role in HCC. We note that *RB1-T* is also overexpressed. However, there are no edges between *RB1-T* and RB1 phosphorylation sites (S3 FigB), suggesting that overexpression of *RB1-T* is not the main source of RB1 hyperphosphorylation. In addition, since unphosphorylated RB1 acts as a tumor suppressor, knocking it down does not seem wise. Many efforts rather target inhibiting its phosphorylation and activating its tumor-suppressive properties [48,50]. Furthermore, Indovina et al. [48] mention Cdk inhibitors as possible therapies which can prevent RB1 phosphorylation. Indeed, Ng et al. [26] found an association of overactive CDK1/CDK2/CDK5 kinases and the phenotype associated with mutations in *TP53*. The central role of phosphorylation of RB1 in all networks suggests that inhibition of Cdk can be beneficial for patients in all clusters.

Many of the edges in discovered networks are absent in the public PPI databases. The edge from *TP53-M* to *LECT2-T* is present in *G*_3_, and *TP53-M* is negatively correlated with *LECT2-T* in this cluster (it is also negatively correlated with *LECT2-T* in *G*_2_, but this edge has a low posterior probability). We note that *LECT2-T* is also downregulated in cluster 2 and cluster 3, but not in cluster 1. The downregulation of *LECT2-T* has been previously associated with a poor prognosis in HCC and mutations in *TP53* [51]. Thus, the discovered link between mutations in *TP53-M* and downregulation of *LECT2-T* is plausible, despite being absent in the STRING database. We further noted that *LECT2-M* has an incoming edge from *TCHH-M* in both *G*_2_ and *G*_3_, while *TCHH* mutations are absent in cluster 1. Both *M* nodes are negatively correlated with *LECT2-T* suggesting that *TCHH-M* contributes in a similar way to the molecular phenotype as *TP53-M*. Heterogeneity is a known issue in identifying cancer subtypes. One implication of shared connections of different mutated genes in the discovered networks is that they affect similar downstream genes and may be targeted by similar therapies.

At the same time, some *M* nodes have opposite effects on the same interaction partners, indicating opposite effects of these corresponding mutated genes on the phenotype. *TP53-M* and *CTNNB1-M* share two common connections: HDAC4_S246 and *KMT2D-T*. In both cases, the mutation status of *CTNNB1* and *TP53* are oppositely correlated with their shared interaction partners. The correlation between *TP53-M* and *KMT2D-T* is positive, while the correlation between *KMT2D-T* and other *M* nodes (shown in S3 FigA) including *CTNNB1* is negative. In pancreatic cancer, low expression of *KMT2D* has been associated with a better prognosis [52]. Moreover, knock-out of *KMT2D* has been shown to attenuate cell proliferation and was suggested as a therapeutic target [53]. Opposite effects of *TP53-M* and *CTNNB1-M* on *KMT2D-T* in cluster 3 suggest that co-occurrence of these mutations may diverge the phenotype from phenotypes where *TP53* and *CTNNB1* do not co-occur. Mutations in *CTNNB1* and *TP53* have been considered mutually exclusive in many studies [54]. However, they co-occur in 10% of all samples in the analyzed dataset. The mutual exclusivity was also challenged by a study presenting a detailed case of *TP53/CTNNB1* co-occurrence in the same tumor [55]. In addition, we observe an interesting pattern of co-occurrence of *TP53* and *CTNNB1* across discovered clusters as four out of five co-occurrence cases fall outside of the *TP53*-dominated cluster 2, which can also hint at possible opposite effects of mutations in *TP53* and *CTNNB1* on the phenotype. Our findings align with another HCC classification based on morphological features of the tumor and gene expression [56]. The analysis by Trobenson et al. [56] indicated that *CTNNB1* and *TP53* were associated with opposite effects on the presence of pseudoglands (a histopathologic feature used for HCC characterization in clinics). In addition, the majority of samples with co-occurring *CTNNB1*/*TP53* mutations ended up in the *CTNNB1* cluster based on the gene expression data. However, *CTNNB1/TP53* mutated tumors were associated with clonal progression, in contrast to tumors harboring only *CTNNB1*.

### 1.5 Hub phosphorylation sites

In studies devoted to PPI network characterization, the number of neighbors (degree) of a node in the network is often used to characterize its biological importance [57,58]. Following this logic, we defined two lists of the most connected nodes in the networks discovered by bnClustOmics. In the first list, we included the top twenty nodes with the largest number of connections that are present with non-zero posterior probabilities in two or all networks (S1 File). Such nodes and their direct neighbors represent the most similar parts between the networks. In the second list, we included all nodes with the largest number of cluster-specific connections (S2 File). Interestingly, the nodes in the first list turned out to be *P* nodes (9 out of 20), *M* nodes (9 out of 20) nodes and *T*-nodes (2 out of 20) while the top nodes of the second list were dominated by *PP* nodes (17 out of 20). Hence, of all omics types, phosphorylation sites appear to have the most different neighborhoods between the clusters. While for *CN*, *T*, and *M* nodes, this can be explained by model structural restrictions, for *P* and *PP* nodes, this finding suggests that differences in the interactome between clusters are more substantial at the phosphoproteome level than at the proteome level.

The list of most differentially connected phosphorylation sites includes MAPK1_T185, CTNND1_S252, and GRB14_S372, which are known to play a role in HCC signaling and affect the regulation of cell cycle, apoptosis, and carcinogenesis (S3 Table). Some of these hub-phosphorylation sites have been found to be important in other cancers than HCC, e.g., ANKRD28_S1011, PRKAA2_S491, and TBXA2R_S331. Our networks suggest that they might also play a role in HCC and are thus candidates for further experiments.

MAPK1 is known to be essential for MAP kinase signaling, which is one of the targets of Sorafenib [59–61], a standard-of-care treatment for advanced HCC. The phosphorylation site MAPK1_T185 is increased in cluster 2 and cluster 3 and has a considerable amount of cluster-specific connections in *G*_2_ (Fig 6E). The phosphorylation of another MAPK1 site, namely Y187, is significantly increased in cluster 3 only. Both phosphorylation sites have many references in the PhosphoSitePlus database, and are known to induce carcinogenesis and alter apoptosis, and are known drug targets. However, MAPK1 is known to be active if both sites are phosphorylated [62]. The increased phosphorylation of both sites is observed only in cluster 3. At the same time, the role of mono-phosphorylated MAPK1 is not fully understood [63]. Sorafenib which inhibits upstream regulators of MAPK1 [64] was given to six patients from the analyzed cohort, three of which were assigned to cluster 2 and three to cluster 3. Five out of six patients had to discontinue treatment due to side-effects, but patients from cluster 3 on average tolerated the therapy longer and survived longer than patients who were treated with Sorafenib in cluster 2 (S9 Appendix). This separation aligns well with our clustering, although it is not possible to make stronger conclusions due to a limited number of biopsies and the short duration of treatment.

One of the MAPK1_T185 interaction partners in *G*_2_ is another hub phosphorylation site, PTPN1_S352, whose phosphorylation is increased in cluster 2 only. *PTPN1* is known to play an important role in many liver diseases; however, it can act both as a tumor suppressor, and oncogene in HCC [65]. Most studies suggest its tumor-suppressive role. However, our analysis indicates that increased phosphorylation of PTPN1_S352 is associated with a poor prognosis and increased phosphorylation of MAPK1_T185 in cluster 2. This connection is confirmed in [66], where *PTPN1* was identified as an oncogene, and its knockdown resulted in attenuated Ras activity and MAPK signaling. We found several inhibitors of PTPN1 in The International Union of Basic and Clinical Pharmacology (IUPHAR) / British Pharmacological Society (BPS) Guide to PHARMACOLOGY [67]. All of them have hypoglycaemic and other anti-diabetic effects. Previous studies already pointed out the anti-tumor properties of diabetes drugs on HCC [68]. We believe that investigating strong individual dependencies in cluster-specific networks coupled with DGE might suggest drug candidates and highlight interactions that are important in the context of different subtypes of HCC.

### 1.6 Discussion

Learning biological networks and cancer subtyping based on multi-omics molecular data are challenging problems, which are traditionally addressed by separate computational methods. In this work, we present bnClustOmics, a tool that tackles both problems simultaneously. Our approach can integrate and cluster multi-omics datasets and learn networks consisting of different types of omics variables, each of which characterizes a patient cluster. In simulation studies, we have shown that bnClustOmics outperforms other clustering approaches due to its ability to detect differences in network structures, while other algorithms mostly lack this ability. A major limitation of our method is the necessity to perform feature selection, which is not straightforward in an unsupervised setting. We suggest using a combination of MOFA and DGE analysis based on our simulation studies, but other ways can also be explored in the future. The package bnClustOmics can be applied to any combination of omics types and is not limited to the five omics types analyzed in this HCC cohort. In the current implementation, there is no possibility to learn the edges between discrete nodes. This feature can further refine clustering, but it makes sense only for larger datasets due to the extreme sparsity of the mutation data.

We applied bnClustOmics to an HCC dataset comprising five different omics types. Similar to previous studies [26,40,56], the three discovered clusters are associated with mutations in *CTNNB1* and *TP53*, and the BCLC stage. Our patient clustering is significantly associated with survival with and without adjustment for the BCLC stage. Cluster 2 is dominated by samples with mutated *TP53* and is associated with a poor prognosis. Samples in which *CTNNB1* and *TP53* co-occur are mostly found in cluster 1 and cluster 3. Moreover, we find that *CTNNB1* and *TP53* have opposite effects on the expression of the transcript *KMT2D* and the phosphorylation site HDAC4_S246 in the learned networks. These findings might explain why *CTNNB1* and *TP53* show mutual exclusivity patterns [69,70] and are associated with opposite effects of the phenotype [56] in some cohorts.

On a more general level, our analysis suggests that the discovered clusters are associated with changes in signaling networks as identified by substantial differences in the neighborhoods of phosphorylation sites. The differences between interactions partners are the largest on the phosphoproteome level, suggesting that this omics type brings a major contribution to the result of the network-based clustering highlighting the importance of phosphoproteome data for further studies.

Cluster-specific networks suggest that hyperphosphorylation of RB1 is associated with mutations in *TP53, CTNNB1*, and *FAT4* but not with overexpression of *RB1* at the transcriptome level. This finding aligns with previous studies suggesting that unphosphorylated RB1 acts as a tumor suppressor, while hyperphosphorylation of RB1 contributes to carcinogenesis [48]. Hence therapies that inhibit phosphorylation of RB1 such as Cdk inhibitors may be a promising treatment strategy.

Overall, our analysis has shown that including associations between different omics types in the clustering model is an important step towards defining cancer subtypes and their molecular makeup comprehensively. These novel associations may improve the selection of effective personalized therapies.

## 2 Methods

### 2.1 Data

We applied bnClustOmics to the HCC data analyzed in [26] (S1 Text). The full dataset comprises 51 biopsies from 49 patients with HCC diagnosis. For each patient, DNA, RNA, proteome, and phosphoproteome data are available. For two patients, two sets of biopsies were available from two genetically different HCC tumors. In addition, we obtained data from 15 biopsies from healthy livers for transcriptome analysis and 11 biopsies for proteome and 10 for phosphoproteome analysis from the same study. A detailed description of sequencing, library preparation, transcript quantification, and SWATH analysis can be found in [26]. We obtained the normalized data from Ng et al. [26] and performed data imputation and batch-correction where applicable (S7 Appendix, S1 Text). One sample was hypermutated with over 9000 mutated genes and was excluded from the analysis. Consequently, we included 50 biopsies from 48 patients in the study.

### 2.2 Bayesian network mixture model

We assume that the data *D* consisting of *N* observations is generated from a mixture of *K* components with weights *τ_k_*. Each component is a Bayesian network 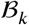, a directed probabilistic graphical model representing a factorization of the joint distribution of the random variables *X*_1_, …, *X_n_*. The random variables are used to model omics features in the analyzed dataset (*M*, *CN*, *T*, *P* and *PP*). Each patient sample *D_i_* represents a vector of *n* values (one for each *X_j_*) and is generated from a model 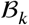, depending on the value of a hidden variable *Z_i_* [14],

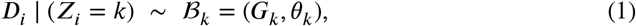

where *G_k_* is a DAG and *θ_k_* are the parameters of the local probability distributions (LPD).

A Bayesian network mixture model was first suggested in [14] for (single-omics) binary mutation data. In our model, each network consists of binary (mutations), ordinal (CNA), and continuous variables (transcriptome, proteome, and phosphoproteome). We denote the set of indices of all binary, ordinal, and continuous nodes by Ω, Φ, and Ψ, respectively. The quantities *n_b_, n_o_*, and *n_c_* are the numbers of binary, ordinal, and continuous random variables, respectively, in the network. We model the LPD for each continuous node *X_ψk_, ψ* ∈ Ψ, of each mixture component by linear regression on its parents in graph *G_k_*,

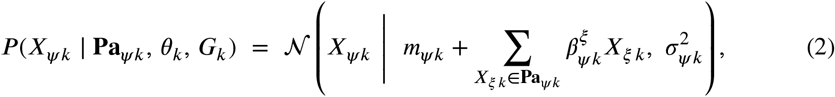

where **Pa**_*ψk*_ is the set of parents of node *X_ψk_* in graph *G_k_*. The set of parameters of the LPDs of continuous nodes includes a vector of regression intercepts *m_k_*, a vector of standard deviations *σ_k_*, and a vector of regression coefficients *B_ψk_* defined for all nodes with non-empty parent set. Given a graph *G_k_*, the Gaussian Bayesian network model above can be equivalently parameterized using a vector of unconditional means *μ_k_* and a covariance matrix ∑_*k*_ (S4 Appendix). We use both parametrizations interchangeably. Binary and ordinal nodes are not allowed to have parents by assumption. For binary nodes *X_ωk_*, we assume that the LPDs are defined by the parameters

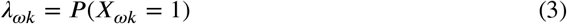

and for ordinal nodes *X_ϕk_*, we use the Gaussian approximation

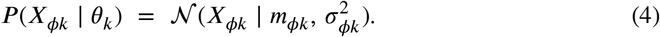

We denote the set of all parameters of a mixture component *k* by *θ_k_* = (*λ_k_, μ_k_*, ∑_*k*_).

### 2.3 EM algorithm

Following [14] we use an EM algorithm for learning Bayesian network mixture models. We denote by *D_i_* the *i*-th observation in the dataset, representing a vector of omics measurements of one patient (or one biopsy in case of multiple biopsies per one patient). The algorithm proceeds as follows:

1. Initialize cluster membership probabilities *γ_ik_* of patient *i* being in cluster *k* (Section 2.7)
2. Given *γ_ik_*, perform MAP structure search and estimate DAGs 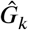 (Section 2.5)
3. Given estimated DAGs 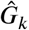, iterate *q* times:
  - (M-step) Compute MAP parameters 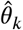 (S5 Appendix)
  - (E-step) Update membership weights

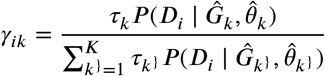

and cluster weights

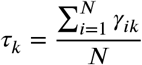 (Section 2.6)
4. Iterate steps 2 and 3 until convergence

The internal cycle with *q* iterations is added for computational efficiency because parameter updates are computationally less expensive than structure search. Hence, for each update of the structures, we perform *q* updates of the parameters. We learn cluster membership assignments for all patients *D_i_* and MAP networks 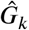. Once the EM algorithm has converged, bnClustOmics can optionally perform sampling from the posterior distribution and the output includes the matrices of estimated probabilities of all edges (Section 2.5).

The main differences to the procedure in [14] are a different set of parameters *θ_k_* and network structural constraints due to the multi-omics extension and differences in data types.

### 2.4 Network score

For assessing how well the network structure fits the data, we use the BGe score [71,72]. In addition to the model assumption specified in Eq 2, the BGe score requires technical assumptions on likelihood and parameter prior [71]. The network score *R*(*G_k_* | *D*) then decomposes over continuous nodes as

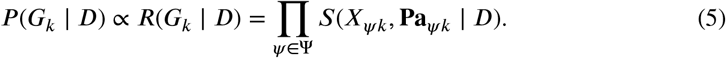

By our model design, nodes *X_ϕ_* and *X_ω_*, corresponding to mutations and copy number changes, are not allowed to have any parents. Hence, the terms *S*(*X_ϕk_*, **Pa***_ϕk_* | *D*) = *S*(*X_ϕk_* | *D*) and *S*(*X_ωk_*, **Pa***_ϕk_* | *D*) = *S*(*X_ωk_* | *D*) are constant for all possible graphs. For this reason, we exclude these terms when performing structure search and the product in Eq 5 runs only over nodes *X_ψk_*. However, nodes *X_ϕk_* and *X_ωk_* may enter the equation as parents of *X_ψk_*.

### 2.5 Structure search

At each step of structure search, we use the iterative order MCMC scheme introduced in [30] and implemented in the R-package BiDAG [31], which proved to be superior to many other methods for MAP structure search in simulation studies [30]. An optional step after the MAP graph has been found is to sample graphs from the posterior distribution using the order MCMC scheme [30]. This step allows us to estimate consensus models by averaging over a sample of *L* graphs from the posterior distribution. In particular, the posterior probability of an edge *e_ξψk_* between nodes *X_ξk_* and *X_ψk_* in the graph *G_k_* is estimated as:

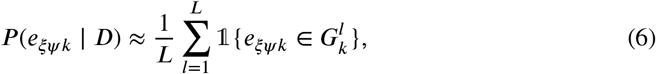

where 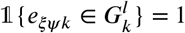 if the edge *e_ξψk_* is present in structure 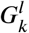 and 0 otherwise. Edges whose posterior probabilities are lower than a defined posterior threshold are excluded from the resulting consensus structure [30].

We use the iterative MAP search at the second step of the EM algorithm and perform sampling once after the EM has converged to compute posterior probabilities of single edges and identify consensus graphs.

To construct graphs for the downstream analysis, we made a list of edges whose posterior *P*(*e_ξψk_* | *D*) is higher than 0.9 for at least one cluster *k* (the threshold was chosen based on our simulation studies). In addition, we selected all edges whose sum of posteriors in all clusters 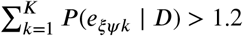, while the threshold for individual networks is lower: *P*(*e_ξψk_* | *D*) > 0.5 for at least one cluster *k*. Finally, we constructed the graphs *G_k_* by including edges from the selected list if their posterior *P*(*e_ξψk_* | *D*) > 0.4. The reason behind this selection process is finding high-confidence cluster-specific interactions while not dismissing similarities at lower (but non-zero) posterior levels.

### 2.6 Cluster membership weights

Updating the membership weights *γ_ik_* requires assessment of the likelihoods 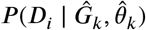. The decomposition provided by the Bayesian network model allows us to integrate discrete and continuous data types in measuring how well an observation *D_i_* (a vector consisting of *n_c_* continuous, *n_o_* ordinal, and *n_b_* binary components) fits a DAG 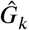 and parameters 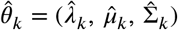:

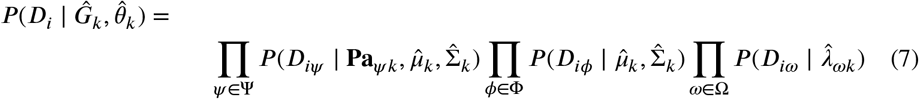

The detailed formulas for computing the likelihoods are given in S6 Appendix. We have extended the R-package BiDAG, such that the function scoreagainstDAG is able to accommodate mixed data.

### 2.7 Starting membership weights

In general, the EM algorithm does not guarantee finding the global maximum, and the local maximum it finds will depend on the starting point. For this reason, we use a non-random starting point in order to start in a parameter region of high likelihood and help mitigate the local optima issue. By default (and for the HCC data), the starting cluster membership of patients is defined via running mclust on the first *K* +2 principal components after applying PCA to the original data. Our simulation studies have shown that dimension reduction via PCA as a starting point improves the results of mclust. The initial membership weights are then defined as

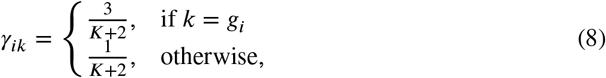

where *g_i_* denotes the cluster assignment of the *i^th^* observation by mclust. PCA is applied only to define the initial membership weights, but the EM algorithm is then applied to original non-reduced data. With a non-random starting point, by default, bnClustOmics runs the EM only once (the results of simulation studies are shown for one run). However, for the HCC dataset, we restarted the EM five times and selected the model with the highest likelihood for each value of *K*.

### 2.8 Allowed edges

By design, bnClustOmics only prohibits incoming edges to discrete nodes. In the HCC data analysis, we added more constraints to obtain more biologically relevant networks. The general flow of the information is directed from the DNA to RNA and (pshospho)protein nodes (S4 Table).

Naturally, we allow all possible edges between *P* and *PP* nodes. We do not allow edges between transcripts because the transcripts do not interact directly. When proteome data is not available, it makes sense to approximate protein-protein interactions with transcript-transcript interactions. However, since we have (phospho)proteome data available, we prefer to explain dependencies with more relevant and interpretable edges between (phospho)proteins and between transcripts and proteins.

### 2.9 Edge penalization matrix

When performing structure search, we use the prior information about interactions between the genes included in the networks, following the methodology described by Kuipers et al. [14]. To do this, we modify the default prior distribution over structures *P*(*G_k_*) and replace it with

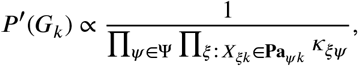

where *κ_ξψ_* defines the penalization factor of the edge *X_ξk_* → *X_ψk_*. Note that *κ_ξψ_* ≥ 1 and these factors do not depend on *k* since prior knowledge does not include cluster assignments. The change of prior leads to replacing of the score terms *S*(*X_ψk_*, **Pa***_ψk_* | *D*) with the terms *S*′(*X_ψk_*, **Pa***_ψk_* | *D*) in Eq 2.4 for all nodes *X_ψk_* with non-empty parent sets:

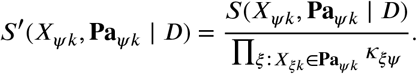

We use the STRING v.11.0 [35] and Omnipath [39] databases to define penalization factors. We penalize the edges by a factor of 2 if they are not found in the databases. The edges corresponding to interactions from the Omnipath database are not penalized. The edges corresponding to the interactions from the STRING database are not penalized if the interaction score is equal to or bigger than 0.5. Otherwise the penalization factor is defined as 2 – 2 * *interaction_score*. In addition, we do not penalize the edges between the same genes of different omics types, e.g., the edges *TP53-T* → *TP53-P* and *TP53-CN* → *TP53-T* are not penalized.

### 2.10 Feature selection

The structure search is the most computationally expensive step of the learning procedure. The complexity of the structure search scheme depends only on the number *n_c_* of continuous nodes in the network (since the product in Eq 5 goes only over continuous nodes) and equals 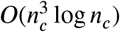 [30]. Hence, for the feasibility of bnClustOmics, we must pre-select the features which we include in the Bayesian networks. Another beneficial point of sensible feature selection is better interpretability since the qualitative analysis is hardly possible for networks with thousands of nodes.

We selected 778 omics features in total (Table 3 in S3 Appendix): 24 *M*, 292 *CN*, 188 *T*, 116 P and 158 *PP*. The main idea behind our feature selection approach was to combine methods that proved to work best in simulation studies (S2 Appendix) with prior knowledge about genes and interactions that are known to be important in HCC signaling (S3 Appendix). In addition to listed criteria we used reasonable filters for selected features: we included only those *M* nodes which are present in at least two samples and *CN* nodes with non-zero variance.

**Table 1.** Distances between cluster centers, *n* = 120. Correspondence between parameters *η, δ* and labels used to define the distances between cluster centers Fig 2A:C (*n* = 120, *n_b_* = 20, *n_c_* = 100). The fourth column represents average *L*_2_ norm between pairs of *μ_i_* and *μ_j_*, *i ≠ j* for all generated mixtures; the range is given in brackets.

| Distance | $\eta$ | $\delta$ | average $L_2$ |
| --- | --- | --- | --- |
| no | 0.1 | 0 | 0.27 (0.03, 0.85) |
| small | 0.2 | 0.1 | 5.78 (4.24, 6.86) |
| medium | 0.3 | 0.2 | 9.44 (7.89, 10.73) |
| large | 0.4 | 0.4 | 14.28 (12.89, 17.37) |

**Table 2.**
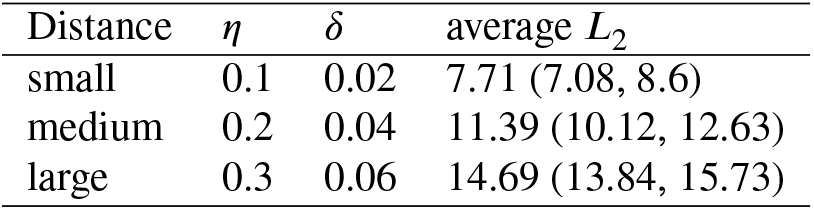
Distances between cluster centers, *n* = 1100. Correspondence between parameters *η*, *δ* and labels used to define the strength of the signal in Fig 2D (*n* = 1100, *n_b_* = 100, *n_c_* = 1000). The fourth column represents average *L*_2_ norm computed pairwise for all *μ_i_* and *μ_j_*, *i ≠ j* for all generated mixtures; the range is given in brackets.

**Table 3.** Summary of feature selection for each omics type. The features are selected as a union of features satisfying the listed criteria. *n_O_* denotes the number of selected features per each omics type.

| omics type | criteria | $n_O$ |
| --- | --- | --- |
| $M$ | MutSigCV ($q < 0.1$)<br>Identified in [36–38] | 24 |
| $CN$ | M, T, P and PP features<br>Identified in [26] | 292 |
| $T$ | MOFA top 50 by absolute weight,<br>P features<br>Targets of P features chosen as TF | 188 |
| $P$ | MOFA top 50 by absolute weight<br>M, T features<br>DE ($q < 0.05$) + present in Omnipath TF database<br>DE ($q < 0.05$) + present in Omnipath KS database | 116 |
| $PP$ | MOFA top 50 by absolute weight<br>DE ($q < 0.05$) + present in Omnipath KS database | 158 |
| all |  | 778 |

### 2.11 Survival analysis

To study the association of clusters with clinical outcomes, we used the Cox proportional hazards model with and without adjustment for clinical stage BCLC. Time was measured in days from the date of diagnosis. In the adjusted model, we excluded BCLC group “0” consisting of one sample, which did not include death events. If two or more biopsies were available for one patient, one of them was included in the analysis if the cluster assignments for all of them were the same. Otherwise, all samples from the patient were excluded. Two samples of patients who were lost-to-followup were considered censored. We used a likelihood ratio test based on the *χ*^2^ distribution to assess the model fit.

### 2.12 Enrichment analysis

Pathway enrichment analysis was performed using the R package ReactomePA [73]. For each omics type, a list of differentially expressed/phosphorylated genes (proteins, phosphoproteins) with FDR adjusted *p*-value smaller than 0.05 was used as input. Pathways enriched with FDR-adjusted *p*-value smaller than 0.05 were selected for visualization.

### 2.13 Differential gene and protein expression analysis

For DGE analysis, we used the R package edgeR [74] for transcriptome data, and limma [75] for proteome and phosphoproteome data. Genes were considered differentially expressed if the FDR-adjusted *p*-value was smaller than 0.05. For variable selection, we compared tumor to healthy samples for all omics types. For the heatmap in Fig 4D, we compared samples in a specific cluster to samples in all other clusters. In the downstream analysis, we have also performed DGE analysis between tumor samples in individual clusters and healthy samples.

## Acknowledgments

We thank Dr. George Rosenberger and Yannick Suter for their expertise, advice, and fruitful discussions on topics presented in this manuscript.

Part of this research was supported by the European Research Council (ERC) Synergy Grant 609883 and SystemsX.ch Research, Technology and Development (RTD) Grant 2013/150.

## Supporting information

### S1 Appendix. Generating Bayesian network mixtures and data in simulations studies

The steps of generating a Bayesian network mixture and the dataset from this mixture include:

1. Generating DAGs *G_k_, k* =1, …, *K*, consisting of *n* variables each, *n_b_* binary and *n_c_* continuous.
  - First, we generate a random DAG *G*_1_ consisting of *n_c_* continuous nodes with the function randomDAG from the R-package pcalg [76]. The parameter prob is set to 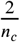 and corresponds to each continuous node *X_ψk_* having one continuous parent on average. Then, the remaining *K* – 1 DAGs in the mixture are generated such that the SHD between each of them and the first structure equals *η*|*E*|, where |*E* | is the number of edges in the first randomly generated structure. We use *η* values in the range 0.1 – 0.4, hence graphs representing different mixture components have a lot of edges in common. From a biological point of view, this makes sense: While some interactions may be altered in a particular cancer subtype, most of them will stay the same (e.g. housekeeping pathways).
  - At the next step, we add random edges from binary to continuous nodes, such that each binary node has 0.5 continuous children on average. These edges model the effects of mutations on other nodes (e.g. transcripts, proteins) and are generated randomly for each mixture component.
2. Generating parameters of local probability distributions (LPDs) for each structure.
  - For 99% of binary nodes *X_ωk_*, frequencies *λ_ωk_* are sampled from beta distribution with parameters *α* = 0.1, *β* = 7, modeling sparse and heterogeneous mutation data. For 1% of binary variables (minimum one variable) frequencies *λ_ωk_* are sampled from a beta distribution with parameters *α* = 0.5 and *β* =1, modeling rare genes, for which higher frequencies are observed in known cancer subtypes.
  - For continuous nodes: regression coefficients *β_ψk_* for nodes with non-empty parent sets are chosen in the range [0.5, 1.5]. Conditional standard deviations *σ_ψk_* are sampled from a normal distribution with mean 0.3 and standard deviation 0.2; to prevent negative values, we use the absolute values of generated numbers.
  - Regression intercepts *m_ψk_* = 0, by default, apart *v* = *δn_c_* nodes *G_k_*. For these *v* nodes, we first sample the *sign* (“+” or with equal probability and then sample randomly in the range [0.5, 1.5] or [−1.5, −0.5]. The parameter *δ* directly impacts how far the centers of distributions *μ_k_* are from each other.
3. Generating data for each mixture component *Z_k_, k* =1, …, *K* using graphs and parameters generated in the previous steps.
  - Generate *N_Z_k__* observations of each binary node *X_ωk_* from a Bernoulli distribution, using parameters *λ_ωk_*.
  - Generate *N_Z_k__* observations of each continuous node *X_ψk_* according to the Eq 2, using parameters *G_k_, m_k_, B_k_, σ_k_*.

We varied two parameters of generated Bayesian network mixtures to see how different algorithms performed depending on the signal strength, defined as *L*2 norm between centers of distributions of mixture components. The first parameter *η* is responsible for the structural difference between networks representing mixture components. The second parameter *δ* was responsible for differences between vectors of regression intercepts *m_k_, k* =1*, …, K*. The tables below represent the correspondence between *x* – *axis* labels used in Fig 2A:D, *η*, *δ* and average *L*_2_ norm of differences between vectors of unconditional means *μ_i_* and *μ_j_*, *i* ≠ *j*.

### S2 Appendix. Feature selection simulation

This simulation study illustrates clustering accuracy of bnClustOmics depending on the selection of relevant features. For simulations we use *n_c_* = 1000 (similarly to the feature selection benchmarking study [77]) and *n_b_* = 100. For clustering, we choose only binary features which equal to 1 in at least one generated sample. In addition we try several approaches to select the continuous features:

1. random selection: 150 continuous features
2. moCluster: 150 continuous features with non-zero loadings defined by sparse consensus PCA
3. MOFA: top 150 features sorted by total absolute weight in all latent factors
4. ranking by the absolute value of normalized mean: top 150 features sorted by 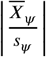, where 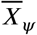 is sample mean and *s_ψ_* is sample standard deviation of *X_ψ_*
5. hybrid of 3. and 4.: mix of MOFA (75 features) and ranking by the absolute value of normalized mean 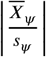 (75 features).

By simulation study design, the default values for all continuous nodes are 0. Hence a ranking by the normalized mean 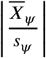 is similar to a ranking by the one-sample *t-test* statistics and defines the variables which most extremely deviate from 0. Feature selection using normalized mean can be seen as a proxy to performing the DGE analysis in the real expression data and selecting genes whose expression mostly deviates from non-tumor samples.

The method moCluster performed best with regard to feature selection in the benchmarking study [77]. However, in that study, the authors generated all variables independently and did not include any interactions in the model. In our simulation, this approach did not perform well. Ranking by normalized mean has shown the best performance. However, MOFA preserves more edges from the original network in subnetworks consisting of selected nodes only. For this reason, when comparing the accuracy of bnClustOmics to other methods in Fig 2D, we show the accuracy of bnClustOmics when the hybrid approach is used for feature selection.

### S3 Appendix. Feature selection for the HCC analysis

To select *M* nodes, we included all mutated genes found significant by MutSigCV tool (*q* < 0.1). In addition, we have added the genes which were found significantly mutated in the TCGA HCC cohort, and the genes identified by HCC studies [36–38] as potential cancer drivers if they were mutated in at least two samples in the HCC dataset.

For nodes of continuous types, we first identified latent factors using MOFA on a subset of features passing standard deviation thresholds (1 for proteome and 2 for transcriptome and phosphoproteome). Five latent factors have been identified by MOFA. Consequently, we selected the top 50 features for each omics type by the total absolute weight of features in all latent factors.

We extended the selected *P* and *PP* features by performing the DGE analysis and picking the differentially expressed features (*q* <0.05), which are also present in the kinase-substrate database Omnipath. Their crucial role in cancer development explains our interest in kinases. Most protein kinases promote cell proliferation, survival, and migration. Furthermore, their aberrant activity is often associated with cancer development [78]. The standard-of-care HCC treatment, Sorafenib, is also a multi-kinase inhibitor.

For *P* nodes, we also selected differentially expressed features present in the transcription factor (TF) database Omnipath, confidence level B.

We have also extended each omics feature set with genes present in selected features of other omics sets for consistency and interpretability of networks. For example, for the possibility of discovering an edge *TP53-M* → *TP53-P*, we have included TP53 at the protein level. The same reasoning stands behind our choice of *CN* nodes, which were selected as a union of gene features selected from transcriptome, proteome, and phosphoproteome. In addition, we included *CN* nodes identified as potential drivers in [26]. We excluded *CN* nodes that had 0 variance in the HCC dataset.

### S4 Appendix. Two ways to parametrize a Gaussian Bayesian network

In this section, we drop indices *k*, so the equations are valid for all mixture components. There are two ways to parametrize a Gaussian Bayesian network. One way is via a vector of regression intercepts *m*, a noise vector *σ*, and regression coefficients *B* = {*β_ψ_*} as in Eq 2. The second way is via a vector of unconditional means *μ* and a covariance matrix ∑. Given the DAG *G*, the parameters {*μ*, ∑} can be transformed into equivalent parameters {*m, B, σ*} [71], where regression coefficients *β_ψ_* are computed only for those nodes, which have non-empty parent sets in a DAG *G*. Let *∑_VW_* be the block of the covariance matrix consisting only of rows with indices *V* and columns with indices *W*. And let *W* be the parents of node *V* in the graph *G*, then

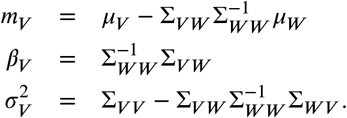

For convenience, we use both parametrizations interchangeably. For example, in defining parameters of simulation studies it is more convenient to use {*m, B, σ*}, while in the description of the EM algorithm we use {*μ*, ∑}.

### S5 Appendix. MAP parameters estimation

Let 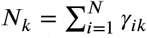. For binary (mutation) nodes, we follow [14] and parametrize local probability distributions as *P*(*X_ωk_* = 1) = *λ_ωk_*, with a beta prior on *λ_ωk_* with hyperparameters 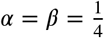. The posterior of *λ* follows a beta distribution as well, so we compute the MAP parameters for all binary nodes *X_ω_* in the M step of the EM algorithm as follows:

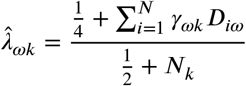

For continuous and ordinal nodes we use the BGe score and assume a normal-inverse-Wishart prior on the parameters *μ* and ∑ [71]:

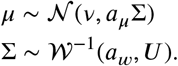

The posterior is then also normal-inverse-Wishart and the MAP parameters are computed as follows:

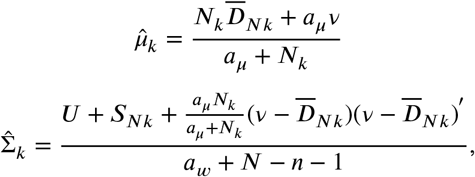

where 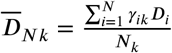 and 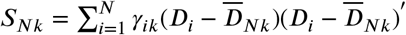. The values of the hyperparameters *a_μ_, a_w_*, the prior mean vector *v* and the parametric matrix *U* by default are set as follows [79]:

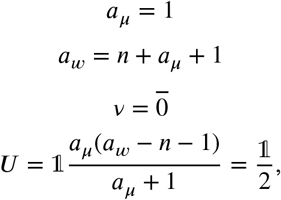

where 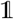 is the identity matrix, and 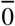 is a vector consisting of zeros.

When MAP graphs *G_k_* and parameters 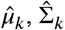 are estimated, the estimates 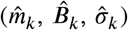 are computed according to S4 Appendix.

### S6 Appendix. Local likelihoods formulas

For binary nodes, we compute likelihoods as

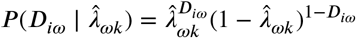

for all binary nodes.

For continuous nodes *X_ψ_*, we compute Gaussian likelihoods according to the model specified in Eq 2,

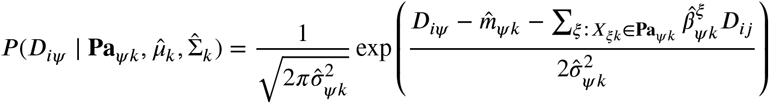

For copy number nodes *X_ϕ_*, we use a similar Gaussian likelihood, but the sum over parents is dropped due to structural assumptions:

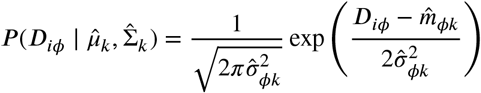

### S7 Appendix. Data pre-processing

For transcriptome data, the pre-processing steps included:

- Like in [26] gene-level expected counts were upper-quartile-normalized to 1000.
- log_2_ transformation.

For proteome data, the pre-processing steps included:

- log_2_ transformation.
- Normalization by median substraction.
- Filtering out proteins which were detected in less than 50% of samples.
- For clustering only: imputation of missing values using the R package impute [80]. For differential expression analysis, we used unimputed values.

For phosphoproteome data, the pre-processing steps included:

- log_2_ transformation.
- Normalization by median substraction.
- Filtering out proteins which were detected in less than 50% of samples.
- Batch correction with the R package edgeR.
- For clustering only: imputation of missing values using the R package impute [80]. For differential expression analysis, we used unimputed values.

The CNA data was obtained at the gene level from the study by Ng et al. [26]. The copy number status was derived from the log-ratio and takes values from 2 to −2, which denote [81]:

- 2: amplification
- 1: copy gain indicates a low-level gain
- 0: copy number neutral
- -1: shallow deletion indicates a shallow loss, possibly a heterozygous deletion
- -2: deep deletion indicates a deep loss, possibly a homozygous deletion

This way, despite the ordinal nature of the CNA data, the range of the values justifies the normal approximation.

### S8 Appendix. High-confidence connections of *TP53-M* in *G*_2_

We investigated the edges outgoing from *TP53-M* whose posterior probabilities in *G*_2_ is higher than 0.9 (p_cl2> 0.9, Table 4). All four identified edges are specific to *G*_2_, as their posteriors in other networks are lower than 0.4. We further considered only gene products that were differentially expressed in cluster 2 (p_DE2< 0.05). Three edges satisfied these criteria. Eventually, we focussed on the edge connecting *TP53-M* and *TERT-T*, since it was also found in the STRING database.

**Table 4.**
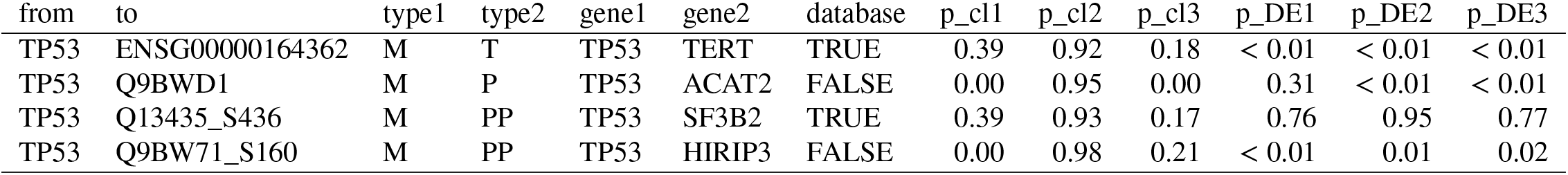
Connections of *TP53-M* node whose posterior probability is larger than 0.9 in network *G*_2_ representing cluster 2. p_cl columns report posterior probabilities for corresponding clusters; p_DE columns report adjusted *p*-values of nodes in column “to” from the DGE analysis.

**Table 5.**
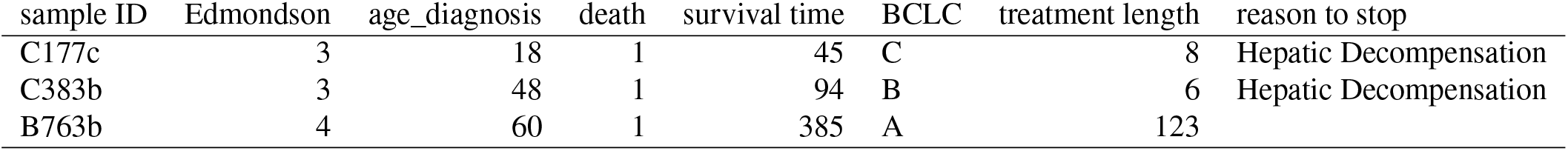
Clinical information about patients who received treatment with Sorafenib and ware assigned to cluster 2 in the clustering by bnClustOmics. Survival time and length of treatment are reported in days.

**Table 6.** Clinical information about patients who received treatment with Sorafenib and were assigned to cluster 3 in the clustering by bnClustOmics. Survival time and length of treatment are reported in days.

| sample ID | Edmondson | age_diagnosis | death | survival time | BCLC | treatment length | reason to stop |
| --- | --- | --- | --- | --- | --- | --- | --- |
| C795 | 2 | 53 | 1 | 625 | A | 58 | Intolerant to side-effects |
| C346b | 2 | 72 | 0 | 1663 | A | 118 | Intolerant to side-effects |
| B983b | 2 | 74 | 1 | 485 | B | 57 | Intolerant to side-effects |

### S9 Appendix. Responses to treatment with Sorafenib

Six out of all 48 patients in the analyzed cohort were treated with Sorafenib. All of these patients were assigned to either cluster 2 or cluster 3 and none to cluster 1. Edmondson grade, survival, and side-effects to Sorafenib were mostly similar within clusters. Two of three patients in cluster 2 experienced hepatic decompensation within several days after the start of treatment and had to stop it. Patients from cluster 3 could, on average, tolerate the side effects longer. All patients in cluster 3 survived longer than patients in cluster 2 even within the same clinical stage.

**S1 Fig.**
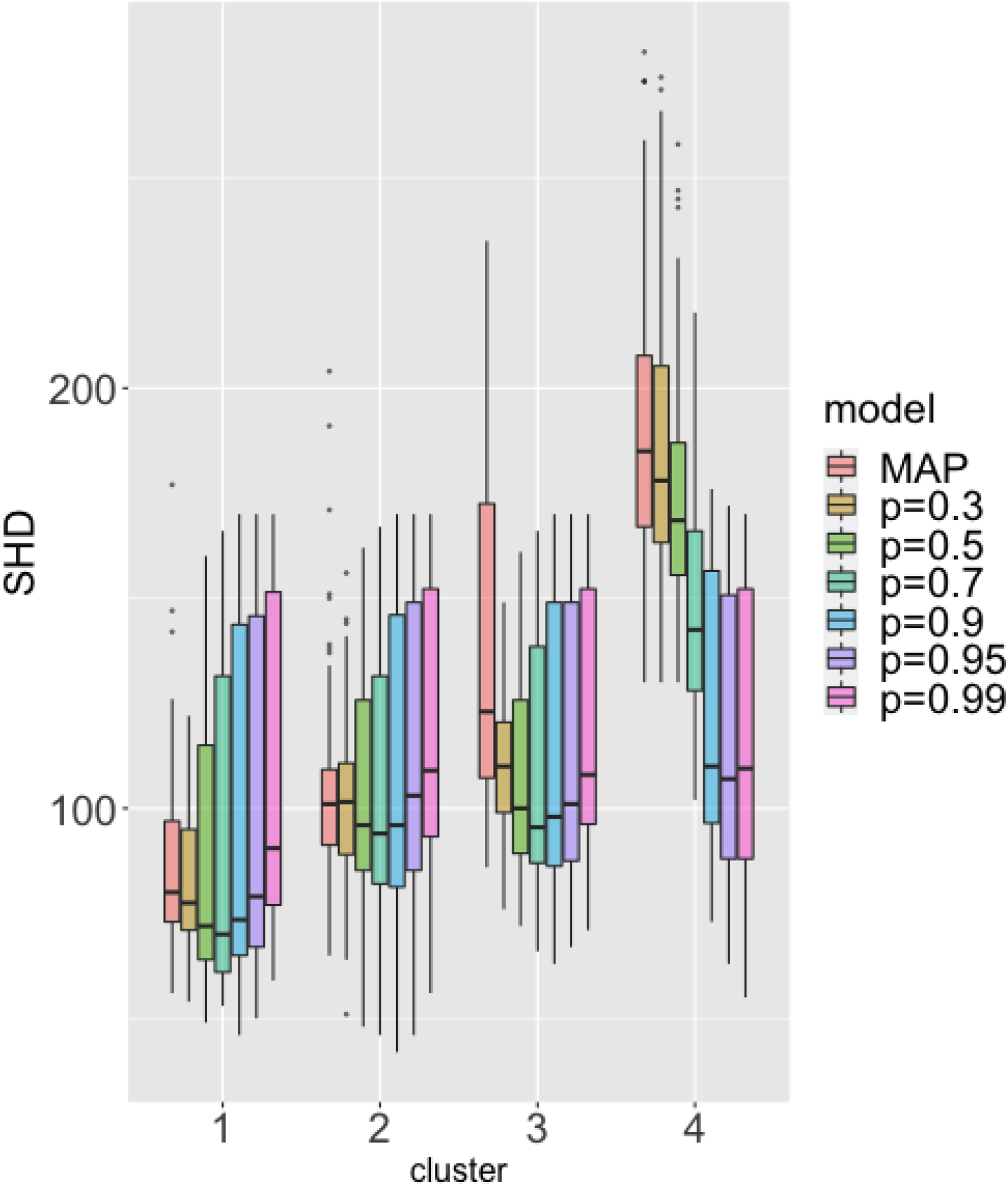
SHD between estimated graphs and the ground truth. 50 BN mixtures were generated with unequal mixture weights: *N*_*Z*_1__ = 150, *N*_*Z*_2__ = 100, *N*_*Z*_3__ = 50, *N*_*Z*_4__ = 20 (cluster 1, cluster 2, cluster 3 and cluster 4). Distance between cluster centers is set to medium. bnClustOmics was used for clustering. The output MAP and consensus structures were compared to the ground truth CPDAG.

**S2 Fig.**
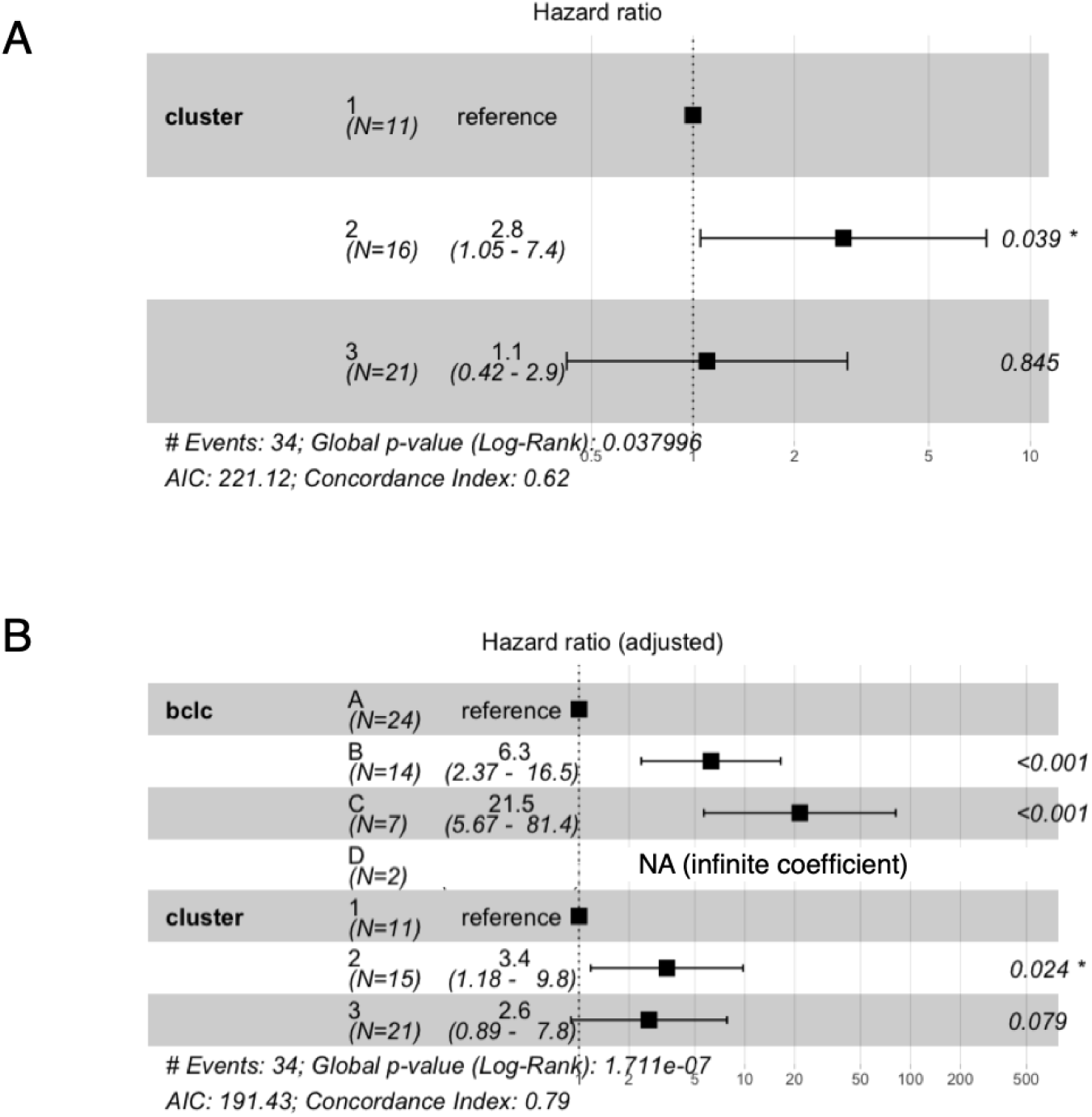
Hazard ratios. Hazard ratios of discovered clusters with (B) and without (A) adjustment for the BCLC stage.

**S3 Fig.**
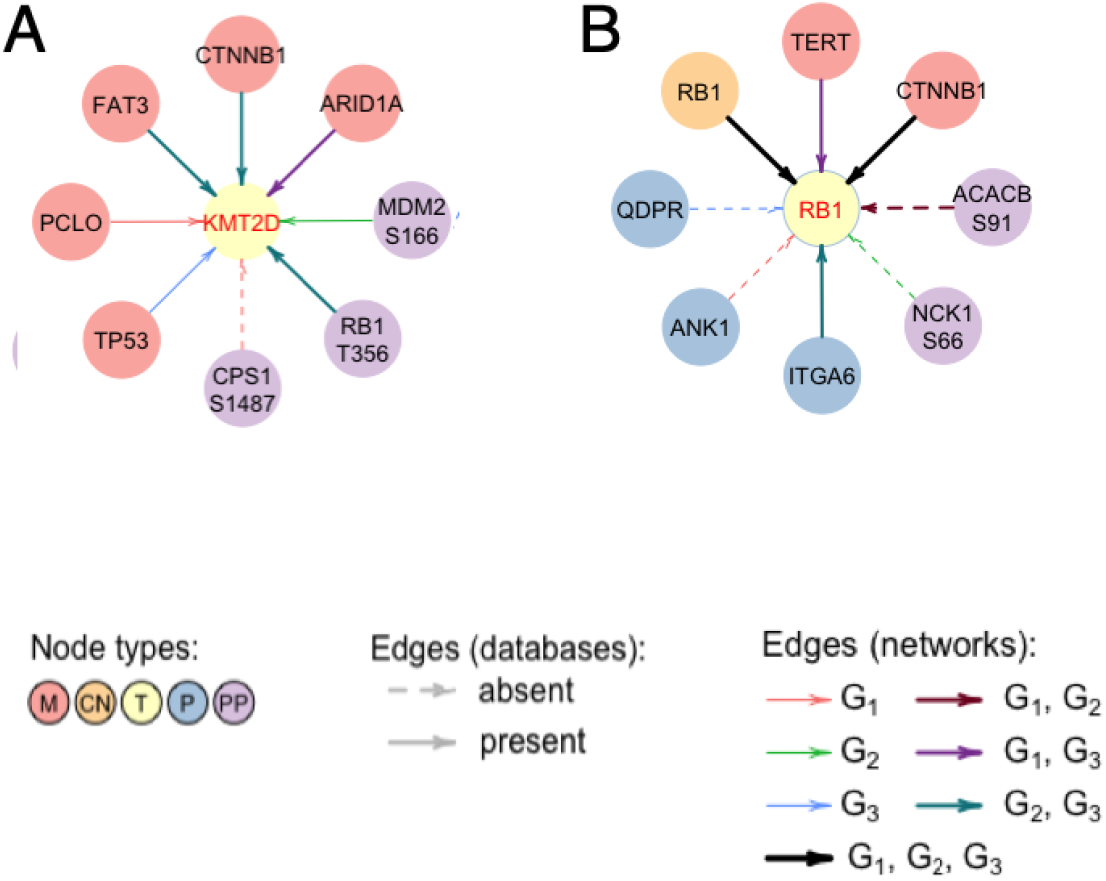
Connections of *KMT2D* and *RB1* transcripts in networks discovered by bnClustOmics.

**S1 Table.** Cox model fit. Summary of the likelihood ratio test for Cox proportional hazards models based on assignments obtained by clustering algorithms. The number of clusters *K* = 3 in all cases. For all algorithms apart from bnClustOmics and MOFA, all available omics features were used as input. For MOFA, standard deviations filters (1 for *P* features, 2 for *T* and *PP*, 0.5 for *CN* features) were applied as recommended by the authors of the method.

| algorithm | $p$-value | $p$-value (BCLC-adjusted) |
| --- | --- | --- |
| mclust | 0.17 | 0.32 |
| hclust | 0.0018 | 0.137 |
| kmeans | 0.024 | 0.77 |
| iClusterPlus | 0.06 | 0.67 |
| CIMLR | 0.94 | 0.39 |
| CIMLR ($T,P,PP$) | 0.017 | 0.20 |
| MOFA | 0.13 | 0.21 |
| bnClustOmics | 0.038 | 0.043 |

**S2 Table.** Signaling pathways enriched with direct interactors of *M* nodes in networks discovered by bnClustOmics. FDR values reflect the enrichment of KEGG signaling pathways with children of *M* nodes in cluster-specific networks. FDR values below 0.05 suggest significant enrichment.

| pathway | $G_1$ | $G_2$ | $G_3$ |
| --- | --- | --- | --- |
| Hepatocellular carcinoma | 0.0007 | <0.0001 | <0.0001 |
| Proteoglycans in cancer | 0.0015 | 0.0007 | <0.0001 |
| PI3K-Akt signaling | 0.0002 | 0.0024 | <0.0001 |
| Cellular senescence | 0.0185 | 0.0039 | 0.00051 |
| wnt signaling | 0.0033 | >0.05 | 0.00058 |
| p53 signaling | 0.0219 | 0.0108 | 0.0079 |
| Insulin signaling | >0.05 | 0.0109 | <0.0001 |
| mTOR signaling | >0.05 | 0.0157 | 0.0031 |
| Cell Cycle | 0.0095 | 0.0387 | 0.0071 |
| AMPK signaling | >0.05 | 0.0387 | 0.0071 |
| HIF-1 signaling | >0.05 | >0.05 | 0.00062 |
| MAPK signaling | >0.05 | >0.05 | 0.0358 |
| JAK-STAT signaling | >0.05 | >0.05 | 0.0166 |
| Rap-1 signaling | >0.05 | >0.05 | 0.0358 |
| Hippo signaling | 0.0033 | >0.05 | >0.05 |
| PD-L1 expression and PD-1 checkpoint pathway in cancer | >0.05 | 0.016 | >0.05 |
| ErbB signaling | >0.05 | >0.05 | <0.0001 |

**S3 Table.** Known functions of most connected phosphorylation sites. A list of phosphorylation sites with more than 15 cluster-specific interaction partners and their known functions in HCC and other cancers according to the PhosphoSitePlus database.

| ID | Gene_ID | DB | In liver studies | In HCC studies | Downstream effects | Upstream regulators | Known treatments |
| --- | --- | --- | --- | --- | --- | --- | --- |
| Q96T23_S1345 | RSF1 | yes | yes | - | - | - | yes |
| O15084_S1011 | ANKRD28 | yes | - | - | intracellular localization | CAMK2D | - |
| Q9UJM3_S273 | ERRFI1 | yes | yes | - | - | EGFR | - |
| Q9Y2T1_S70 | AXIN2 | yes | - | - | - | - | - |
| B1AK53_S642 | ESPN | - | - | - | - | - | - |
| P00533_S1166 | EGFR | yes | yes | yes | activates signalling cascades | CAMK2A | yes |
| P28482_T185 | MAPK1 | yes | yes | yes | altered apoptosis cell growth | - | yes |
| P05556_S785 | ITGB1 | yes | - | yes | cytoskeletal reorganization | PKCA | yes |
| P21333_S1734 | FLNA | yes | - | - | - | - | - |
| O14828_S76 | SCAMP3 | yes | - | yes | - | - | yes |
| O60716_S252 | CTNND1 | yes | yes | yes | carcinogenesis induced | - | yes |
| P18031_S352 | PTPN1 | yes | - | - | - | - | - |
| P19878_S312 | NCF2 | yes | - | - | - | - | - |
| P21731_S331 | TBXA2R | yes | - | - | signalling pathway regulation | PKACA | yes |
| P54646_S491 | PRKAA2 | yes | - | - | altered autophagy | - | yes |
| P62753_S244 | RPS6 | yes | - | yes | - | mTOR TSC2 | yes |
| Q13557_T287 | CAMK2D | yes | yes | yes | - | - | yes |
| Q14449_S372 | GRB14 | yes | - | yes | cell cycle regulation | - | yes |
| Q8IXS6_S228 | PALM2 | yes | - | - | - | - | - |

**S4 Table.** Allowed edges between features in the HCC analysis. Allowed edges (i.e., not blacklisted) between and within omics types in the HCC analysis. Let X and Y denote gene names. Then, all edges from *CN* nodes to *P* nodes of the same genes are encoded as from X-*CN* to X-*P*. Edges between any two genes are encoded as edges between X-*CN* and Y-*P* (this includes the case when X equals Y).

| from | to | example of biological interpretation |
| --- | --- | --- |
| *X-M* | *Y-T, Y-P, Y-PP* | functional interaction |
| *X-CN* | *X-T, X-P, X-PP* | central dogma of molecular biology |
| *X-T* | *X-P, X-PP* | central dogma of molecular biology |
| *X-P* | *Y-T* | transcription factor and its target, functional interaction |
| *X-PP* | *Y-T* | transcription factor and its target, functional interaction |
| *X-P, X-PP* | *Y-P, Y-PP* | physical interaction, functional interaction |

The biological interpretation of most edges is straightforward, however, edges of type X-*M* → Y-*P/T/PP*, for two genes X and Y, are rarely considered when learning the networks. Mehnert et al. [21] have shown via experiments that a cancerous mutation in gene X can change the interactome of its protein product X-*P* without affecting the expression of X-*P* itself. Say Y-*P* is one of the interactors of X-*P* affected by a mutation in gene X. In this case, we can observe a statistical dependency between a mutation node X-*M* and a protein node Y-*P* (but not between X-*P* and Y-*P*). Such dependencies are particularly interesting because they help understand the links between genotypes and phenotypes.

## S1 Text. Data and code availability

The sequencing datasets are available at European Genome-phenome Archive under accessions EGAS00001005073 (whole-exome sequencing), EGAS00001005074 (RNA-sequencing). The mass spectrometry proteomics and phospho-proteomics data have been deposited to the ProteomeXchange Consortium via the PRIDE partner repository under accessions PXD025705 and PXD025836.

The pre-processed reduced and non-reduced multi-omics datasets used for clustering by bnClustOmics and other methods in this study as well as the code and results of simulation studies and HCC clustering are available at the GitHub repository https://github.com/cbg-ethz/HCC. Patient IDs were encrypted.

The R-package bnClustOmics is available at the GitHub repository https://github.com/cbg-ethz/bnclustOmics.

**S1 File. Top twenty most similarly connected nodes and their interactions partners in cluster-specific networks.**

**S2 File. Top twenty most differently connected nodes and their interactions partners in cluster-specific networks.**

## References

1. Wu Y, Liu Z, Xu X. Molecular subtyping of hepatocellular carcinoma: A step toward precision medicine. Cancer Communications. 2020;40(12):681–693. doi:10.1002/cac2.12115.

2. Cai M, Li L. Subtype identification from heterogeneous TCGA datasets on a genomic scale by multi-view clustering with enhanced consensus. BMC Medical Genomics. 2017;10(S4). doi:10.1186/s12920-017-0306-x.

3. Kamoun A, Cancel-Tassin G, Fromont G, Elarouci N, Armenoult L, Ayadi M, et al. Comprehensive molecular classification of localized prostate adenocarcinoma reveals a tumour subtype predictive of non-aggressive disease. Annals of Oncology. 2018;29(8):1814–1821. doi:10.1093/annonc/mdy224.

4. Jiang YZ, Liu Y, Xiao Y, Hu X, Jiang L, Zuo WJ, et al. Molecular subtyping and genomic profiling expand precision medicine in refractory metastatic triple-negative breast cancer: the FUTURE trial. Cell Research. 2020;31(2):178–186. doi:10.1038/s41422-020-0375-9.

5. Nutt CL, Mani DR, Betensky RA, Tamayo P, Cairncross JG, Ladd C, et al. Gene expression-based classification of malignant gliomas correlates better with survival than histological classification. Cancer Res. 2003;63(7):1602–1607.

6. Pierre-Jean M, Deleuze JF, Floch EL, Mauger F. Clustering and variable selection evaluation of 13 unsupervised methods for multi-omics data integration. Briefings in Bioinformatics. 2019;21(6):2011–2030. doi:10.1093/bib/bbz138.

7. Rappoport N, Shamir R. Multi-omic and multi-view clustering algorithms: review and cancer benchmark. Nucleic Acids Research. 2018;46(20):10546–10562. doi:10.1093/nar/gky889.

8. Tini G, Marchetti L, Priami C, Scott-Boyer MP. Multi-omics integration—a comparison of unsupervised clustering methodologies. Briefings in Bioinformatics. 2017;20(4):1269–1279. doi:10.1093/bib/bbx167.

9. Wang D, Gu J. Integrative clustering methods of multi-omics data for molecule-based cancer classifications. Quantitative Biology. 2016;4(1):58–67. doi:10.1007/s40484-016-0063-4.

10. Dimitrakopoulos C, Hindupur SK, Häfliger L, Behr J, Montazeri H, Hall MN, et al. Network-based integration of multi-omics data for prioritizing cancer genes. Bioinformatics. 2018;34(14):2441–2448. doi:10.1093/bioinformatics/bty148.

11. Silverbush D, Cristea S, Yanovich-Arad G, Geiger T, Beerenwinkel N, Sharan R. Simultaneous Integration of Multi-omics Data Improves the Identification of Cancer Driver Modules. Cell Systems. 2019;8(5):456–466.e5. doi:10.1016/j.cels.2019.04.005.

12. Lu M, Zhan X. The crucial role of multiomic approach in cancer research and clinically relevant outcomes. EPMA Journal. 2018;9(1):77–102. doi:10.1007/s13167-018-0128-8.

13. Cajal SR, Sesé M, Capdevila C, Aasen T, Mattos-Arruda LD, Diaz-Cano SJ, et al. Clinical implications of intratumor heterogeneity: challenges and opportunities. Journal of Molecular Medicine. 2020;98(2):161–177. doi:10.1007/s00109-020-01874-2.

14. Kuipers J, Thurnherr T, Moffa G, Suter P, Behr J, Goosen R, et al. Mutational interactions define novel cancer subgroups. Nature Communications. 2018;9(1). doi:10.1038/s41467-018-06867-x.

15. Hofree M, Shen JP, Carter H, Gross A, Ideker T. Network-based stratification of tumor mutations. Nature Methods. 2013;10(11):1108–1115. doi:10.1038/nmeth.2651.

16. Koh HWL, Fermin D, Vogel C, Choi KP, Ewing RM, Choi H. iOmicsPASS: network-based integration of multiomics data for predictive subnetwork discovery. npj Systems Biology and Applications. 2019;5(1). doi:10.1038/s41540-019-0099-y.

17. Vaske CJ, Benz SC, Sanborn JZ, Earl D, Szeto C, Zhu J, et al. Inference of patient-specific pathway activities from multi-dimensional cancer genomics data using PARADIGM. Bioinformatics. 2010;26(12):i237–i245. doi:10.1093/bioinformatics/btq182.

18. Lazareva O, Canzar S, Yuan K, Baumbach J, Blumenthal DB, Tieri P, et al. BiCoN: network-constrained biclustering of patients and omics data. Bioinformatics. 2020;37(16):2398–2404. doi:10.1093/bioinformatics/btaa1076.

19. Grzegorczyk M, Aderhold A, Husmeier D. Overview and Evaluation of Recent Methods for Statistical Inference of Gene Regulatory Networks from Time Series Data. In: Gene Regulatory Networks. Springer New York; 2018. p. 49–94. Available from: https://doi.org/10.1007/978-1-4939-8882-2_3.

20. Xing L, Guo M, Liu X, Wang C, Wang L, Zhang Y. An improved Bayesian network method for reconstructing gene regulatory network based on candidate auto selection. BMC Genomics. 2017;18(S9). doi:10.1186/s12864-017-4228-y.

21. Mehnert M, Ciuffa R, Frommelt F, Uliana F, van Drogen A, Ruminski K, et al. Multi-layered proteomic analyses decode compositional and functional effects of cancer mutations on kinase complexes. Nature Communications. 2020;11(1). doi:10.1038/s41467-020-17387-y.

22. Duan R, Gao L, Gao Y, Hu Y, Xu H, Huang M, et al. Evaluation and comparison of multi-omics data integration methods for cancer subtyping. PLOS Computational Biology. 2021;17(8):e1009224. doi:10.1371/journal.pcbi.1009224.

23. Mo Q, Wang S, Seshan VE, Olshen AB, Schultz N, Sander C, et al. Pattern discovery and cancer gene identification in integrated cancer genomic data. Proceedings of the National Academy of Sciences. 2013;110(11):4245–4250. doi:10.1073/pnas.1208949110.

24. Ramazzotti D, Lal A, Wang B, Batzoglou S, Sidow A. Multi-omic tumor data reveal diversity of molecular mechanisms that correlate with survival. Nature Communications. 2018;9(1). doi:10.1038/s41467-018-06921-8.

25. Argelaguet R, Velten B, Arnol D, Dietrich S, Zenz T, Marioni JC, et al. Multi-Omics Factor Analysis—a framework for unsupervised integration of multi-omics data sets. Molecular Systems Biology. 2018;14(6). doi:10.15252/msb.20178124.

26. Ng CKY, Dazert E, Boldanova T, Coto-Llerena M, Nuciforo S, Ercan C, et al. Proteogenomic characterization of hepatocellular carcinoma. bioRxiv. 2021;doi:10.1101/2021.03.05.434147.

27. Craig AJ, von Felden J, Garcia-Lezana T, Sarcognato S, Villanueva A. Tumour evolution in hepatocellular carcinoma. Nature Reviews Gastroenterology & Hepatology. 2019;17(3):139–152. doi:10.1038/s41575-019-0229-4.

28. Szklarczyk D, Gable AL, Nastou KC, Lyon D, Kirsch R, Pyysalo S, et al. The STRING database in 2021: customizable protein–protein networks, and functional characterization of user-uploaded gene/measurement sets. Nucleic Acids Research. 2020;49(D1):D605–D612. doi:10.1093/nar/gkaa1074.

29. Cobb M. 60 years ago, Francis Crick changed the logic of biology. PLOS Biology. 2017;15(9):e2003243. doi:10.1371/journal.pbio.2003243.

30. Kuipers J, Suter P, Moffa G. Efficient Sampling and Structure Learning of Bayesian Networks. arXiv:180307859v3. 2020;.

31. Suter P, Kuipers J, Moffa G, Beerenwinkel N. Bayesian structure learning and sampling of Bayesian networks with the R package BiDAG. arXiv:210500488. 2021;.

32. R Core Team. R: A Language and Environment for Statistical Computing; 2013. Available from: http://www.R-project.org/.

33. Scrucca L, Fop M, Murphy TB, Raftery AE. mclust 5: clustering, classification and density estimation using Gaussian finite mixture models. The R Journal. 2016;8(1):289–317.

34. Hubert L, Arabie P. Comparing partitions. Journal of Classification. 1985;2(1):193–218. doi:10.1007/bf01908075.

35. Szklarczyk D, Gable AL, Lyon D, Junge A, Wyder S, Huerta-Cepas J, et al. STRING v11: protein–protein association networks with increased coverage, supporting functional discovery in genome-wide experimental datasets. Nucleic Acids Research. 2018;47(D1):D607–D613. doi:10.1093/nar/gky1131.

36. Rao CV, Asch AS, Yamada HY. Frequently mutated genes/pathways and genomic instability as prevention targets in liver cancer. Carcinogenesis. 2016;38(1):2–11. doi:10.1093/carcin/bgw118.

37. Zhang Y, Qiu Z, Wei L, Tang R, Lian B, Zhao Y, et al. Integrated Analysis of Mutation Data from Various Sources Identifies Key Genes and Signaling Pathways in Hepatocellular Carcinoma. PLoS ONE. 2014;9(7):e100854. doi:10.1371/journal.pone.0100854.

38. Kong F, Kong D, Yang X, Yuan D, Zhang N, Hua X, et al. Integrative analysis of highly mutated genes in hepatitis B virus-related hepatic carcinoma. Cancer Medicine. 2020;9(7):2462–2479. doi:10.1002/cam4.2903.

39. Türei D, Korcsmáros T, Saez-Rodriguez J. OmniPath: guidelines and gateway for literature-curated signaling pathway resources. Nature Methods. 2016;13(12):966–967. doi:10.1038/nmeth.4077.

40. Bidkhori G, Benfeitas R, Klevstig M, Zhang C, Nielsen J, Uhlen M, et al. Metabolic network-based stratification of hepatocellular carcinoma reveals three distinct tumor subtypes. Proceedings of the National Academy of Sciences. 2018;115(50):E11874–E11883. doi:10.1073/pnas.1807305115.

41. Maleki F, Ovens K, Hogan DJ, Kusalik AJ. Gene Set Analysis: Challenges, Opportunities, and Future Research. Frontiers in Genetics. 2020;11. doi:10.3389/fgene.2020.00654.

42. Sun X, Wang SC, Wei Y, Luo X, Jia Y, Li L, et al. Arid1a Has Context-Dependent Oncogenic and Tumor Suppressor Functions in Liver Cancer. Cancer Cell. 2018;33(1):151–152. doi:10.1016/j.ccell.2017.12.011.

43. Javanmard D, Najafi M, Babaei MR, Niya MHK, Esghaei M, Panahi M, et al. Investigation of CTNNB1 gene mutations and expression in hepatocellular carcinoma and cirrhosis in association with hepatitis B virus infection. Infectious Agents and Cancer. 2020;15(1). doi:10.1186/s13027-020-00297-5.

44. Lachenmayer A, Alsinet C, Savic R, Cabellos L, Toffanin S, Hoshida Y, et al. Wnt-Pathway Activation in Two Molecular Classes of Hepatocellular Carcinoma and Experimental Modulation by Sorafenib. Clinical Cancer Research. 2012;18(18):4997–5007. doi:10.1158/1078-0432.ccr-11-2322.

45. de Galarreta MR, Bresnahan E, Molina-Sánchez P, Lindblad KE, Maier B, Sia D, et al. *β*-Catenin Activation Promotes Immune Escape and Resistance to Anti–PD-1 Therapy in Hepatocellular Carcinoma. Cancer Discovery. 2019;9(8):1124–1141. doi:10.1158/2159-8290.cd-19-0074.

46. Mertins P,, Mani DR, Ruggles KV, Gillette MA, Clauser KR, et al. Proteogenomics connects somatic mutations to signalling in breast cancer. Nature. 2016;534(7605):55–62. doi:10.1038/nature18003.

47. Zhou X, Lu J, Zhu H. Correlation between the expression of hTERT gene and the clinicopathological characteristics of hepatocellular carcinoma. Oncology Letters. 2015;11(1):111–115. doi:10.3892/ol.2015.3892.

48. Indovina P, Pentimalli F, Casini N, Vocca I, Giordano A. RB1 dual role in proliferation and apoptosis: Cell fate control and implications for cancer therapy. Oncotarget. 2015;6(20):17873–17890. doi:10.18632/oncotarget.4286.

49. Hornbeck PV, Zhang B, Murray B, Kornhauser JM, Latham V, Skrzypek E. PhosphoSitePlus, 2014: mutations, PTMs and recalibrations. Nucleic Acids Research. 2014;43(D1):D512–D520. doi:10.1093/nar/gku1267.

50. Knudsen ES, Wang JYJ. Targeting the RB-pathway in Cancer Therapy. Clinical Cancer Research. 2010;16(4):1094–1099. doi:10.1158/1078-0432.ccr-09-0787.

51. Yang C, Ho M, Chen C, Hsu H, Lee P, Kuo M. The prognostic value of the downregulation of leukocyte cell-derived chemotaxin 2 gene of hepatocellular carcinoma. Journal of Clinical Oncology. 2011;29(15_suppl):10559–10559. doi:10.1200/jco.2011.29.15_suppl.10559.

52. Dawkins JBN, Wang J, Maniati E, Heward JA, Koniali L, Kocher HM, et al. Reduced Expression of Histone Methyltransferases KMT2C and KMT2D Correlates with Improved Outcome in Pancreatic Ductal Adenocarcinoma. Cancer Research. 2016;76(16):4861–4871. doi:10.1158/0008-5472.can-16-0481.

53. Guo C, Chen LH, Huang Y, Chang CC, Wang P, Pirozzi CJ, et al. KMT2D maintains neoplastic cell proliferation and global histone H3 lysine 4 monomethylation. Oncotarget. 2013;4(11):2144–2153. doi:10.18632/oncotarget.1555.

54. Tornesello ML, Buonaguro L, Tatangelo F, Botti G, Izzo F, Buonaguro FM. Mutations in TP53, CTNNB1 and PIK3CA genes in hepatocellular carcinoma associated with hepatitis B and hepatitis C virus infections. Genomics. 2013;102(2):74–83. doi:10.1016/j.ygeno.2013.04.001.

55. Friemel J, Rechsteiner M, Bawohl M, Frick L, Müllhaupt B, Lesurtel M, et al. Liver cancer with concomitant TP53 and CTNNB1 mutations: a case report. BMC Clinical Pathology. 2016;16(1). doi:10.1186/s12907-016-0029-5.

56. Torbenson M, McCabe CE, O’Brien DR, Yin J, Bainter T, Tran NH, et al. Morphological heterogeneity in beta-catenin mutated hepatocellular carcinomas: implications for tumor molecular classification. Human Pathology. 2021;doi:10.1016/j.humpath.2021.09.009.

57. He X, Zhang J. Why Do Hubs Tend to Be Essential in Protein Networks? PLOS Genetics. 2006;2(6):e88. doi:10.1371/journal.pgen.0020088.

58. Goymer P. Why do we need hubs? Nature Reviews Genetics. 2008;9(9):651–651. doi:10.1038/nrg2450.

59. Marisi G, Cucchetti A, Ulivi P, Canale M, Cabibbo G, Solaini L, et al. Ten years of sorafenib in hepatocellular carcinoma: Are there any predictive and/or prognostic markers? World Journal of Gastroenterology. 2018;24(36):4152–4163. doi:10.3748/wjg.v24.i36.4152.

60. Keswani RN, Chumsangsri A, Mustafi R, Delgado J, Cohen EEW, Bissonnette M. Sorafenib inhibits MAPK-mediated proliferation in a Barrett’s esophageal adenocarcinoma cell line. Diseases of the Esophagus. 2008;21(6):514–521. doi:10.1111/j.1442-2050.2007.00799.x.

61. Gedaly R, Angulo P, Hundley J, Daily MF, Chen C, Evers BM. PKI-587 and Sorafenib Targeting PI3K/AKT/mTOR and Ras/Raf/MAPK Pathways Synergistically Inhibit HCC Cell Proliferation. Journal of Surgical Research. 2012;176(2):542–548. doi:10.1016/j.jss.2011.10.045.

62. Pimienta G, Pascual J. Canonical and Alternative MAPK Signaling. Cell Cycle. 2007;6(21):2628–2632. doi:10.4161/cc.6.21.4930.

63. Vázquez B, Soto T, del Dedo JE, Franco A, Vicente J, Hidalgo E, et al. Distinct biological activity of threonine monophosphorylated MAPK isoforms during the stress response in fission yeast. Cellular Signalling. 2015;27(12):2534–2542. doi:10.1016/j.cellsig.2015.09.017.

64. Gong L, Giacomini MM, Giacomini C, Maitland ML, Altman RB, Klein TE. PharmGKB summary. Pharmacogenetics and Genomics. 2017;27(6):240–246. doi:10.1097/fpc.0000000000000279.

65. Huang Y, Zhang Y, Ge L, Lin Y, Kwok H. The Roles of Protein Tyrosine Phosphatases in Hepatocellular Carcinoma. Cancers. 2018;10(3):82. doi:10.3390/cancers10030082.

66. Dubé N, Cheng A, Tremblay ML. The role of protein tyrosine phosphatase 1B in Ras signaling. Proceedings of the National Academy of Sciences. 2004;101(7):1834–1839. doi:10.1073/pnas.0304242101.

67. Protein tyrosine phosphatases non-receptor type (PTPN): protein tyrosine phosphatase non-receptor type 1.;. http://www.guidetopharmacology.org/GRAC/ObjectDisplayForward?objectId=2976.

68. Miyoshi H, Kato K, Iwama H, Maeda E, Sakamoto T, Fujita K, et al. Effect of the anti-diabetic drug metformin in hepatocellular carcinoma in vitro and in vivo. International Journal of Oncology. 2014;45(1):322–332. doi:10.3892/ijo.2014.2419.

69. Kancherla V, Abdullazade S, Matter MS, Lanzafame M, Quagliata L, Roma G, et al. Genomic Analysis Revealed New Oncogenic Signatures in TP53-Mutant Hepatocellular Carcinoma. Frontiers in Genetics. 2018;9. doi:10.3389/fgene.2018.00002.

70. Fujimoto A, Totoki Y, Abe T, Boroevich KA, Hosoda F, Nguyen HH, et al. Whole-genome sequencing of liver cancers identifies etiological influences on mutation patterns and recurrent mutations in chromatin regulators. Nature Genetics. 2012;44(7):760–764. doi:10.1038/ng.2291.

71. Geiger D, Heckerman D. Parameter priors for directed acyclic graphical models and the characterization of several probability distributions. The Annals of Statistics. 2002;30(5). doi:10.1214/aos/1035844981.

72. Kuipers J, Moffa G, Heckerman D. Addendum on the scoring of Gaussian directed acyclic graphical models. The Annals of Statistics. 2014;42(4). doi:10.1214/14-aos1217.

73. Yu G, He QY. ReactomePA: an R/Bioconductor package for reactome pathway analysis and visualization. Molecular BioSystems. 2016;12(2):477–479. doi:10.1039/c5mb00663e.

74. Robinson MD, McCarthy DJ, Smyth GK. edgeR: a Bioconductor package for differential expression analysis of digital gene expression data. Bioinformatics. 2009;26(1):139–140. doi:10.1093/bioinformatics/btp616.

75. Ritchie ME, Phipson B, Wu D, Hu Y, Law CW, Shi W, et al. limma powers differential expression analyses for RNA-sequencing and microarray studies. Nucleic Acids Research. 2015;43(7):e47–e47. doi:10.1093/nar/gkv007.

76. Kalisch M, Mächler M, Colombo D, Maathuis MH, Bühlmann P. Causal Inference Using Graphical Models with the R Package pcalg. Journal of Statistical Software. 2012;47(11). doi:10.18637/jss.v047.i11.

77. Meng C, Helm D, Frejno M, Kuster B. moCluster: Identifying Joint Patterns Across Multiple Omics Data Sets. Journal of Proteome Research. 2015;15(3):755–765. doi:10.1021/acs.jproteome.5b00824.

78. Bhullar KS, Lagarón NO, McGowan EM, Parmar I, Jha A, Hubbard BP, et al. Kinase-targeted cancer therapies: progress, challenges and future directions. Molecular Cancer. 2018;17(1). doi:10.1186/s12943-018-0804-2.

79. Grzegorczyk M. An Introduction to Gaussian Bayesian Networks. In: Systems Biology in Drug Discovery and Development. Humana Press; 2010. p. 121–147. Available from: https://doi.org/10.1007/978-1-60761-800-3_6.

80. Trevor Hastie RT. impute; 2017. Available from: https://bioconductor.org/packages/impute.

81. Cerami E, Gao J, Dogrusoz U, Gross BE, Sumer SO, Aksoy BA, et al. The cBio Cancer Genomics Portal: An Open Platform for Exploring Multidimensional Cancer Genomics Data. Cancer Discovery. 2012;2(5):401–404. doi:10.1158/2159-8290.cd-12-0095.

